# Emergent survival and extinction of species within gut bacterial communities

**DOI:** 10.1101/2024.04.29.591619

**Authors:** Naomi Iris van den Berg, Melanie Tramontano, Rui Guan, Sergej Andrejev, Sonja Blasche, Yongkyu Kim, Martina Klünemann, Ana Rita Brochado, Lajos Kalmar, Anja Telzerow, Peer Bork, Daniel C. Sevin, Athanasios Typas, Kiran R. Patil

**Author notes:** Correspondence to: PB,; NT,; and KRP. These authors contributed equally (Naomi Iris van den Berg, Melanie Tramontano).

## Abstract

Synthetic communities can help uncover metabolic forces shaping microbial ecosystems. Yet, in case of the gut microbiota, culturing in undefined media has prevented detection of metabolic dependencies. Here we show, using chemically defined media, how species survival is jointly determined by supplied resources and community metabolism. We used 63 representative gut bacterial strains and varied inoculum compositions to assemble stable communities in 14 defined media. Over 95% of the species showed markedly improved or diminished performance relative to monoculture in at least one condition, including 153 cases (21%) of emergent survival, i.e., species incapable of surviving on their own but thriving in a community, and 252 (35%) community-driven extinctions. Through single species additions and exclusions, metabolomic analysis, and ecological modelling, we demonstrate how inter-species dependencies – especially in poor media – are mediated by biotic nutrient supply. Our results highlight communal metabolic dividend as a key biotic force promoting emergent survival and diversity.

---

The phylogenetic and metabolic diversity of the gut microbiota is fundamental to the beneficial impact on host through its emergent properties, including resilience to biotic and abiotic perturbations ^1–3^. The diversity of the microbiota is influenced by various factors, with diet being a major determinant contributing substantially more than host genetics ^4–6^. Yet, while diet is closely linked to nutritional requirements of the community members, mechanistic insights remain obscure due to the complex and undefined nature of the human diet and yet limited understanding of the species- and community-scale metabolism of most gut bacteria. We observe a striking discrepancy between growth fitness of a species in monoculture and its relative abundance in the gut microbiota (Fig. 1a), suggesting a crucial role of inter-species interactions.

**Figure 1.**
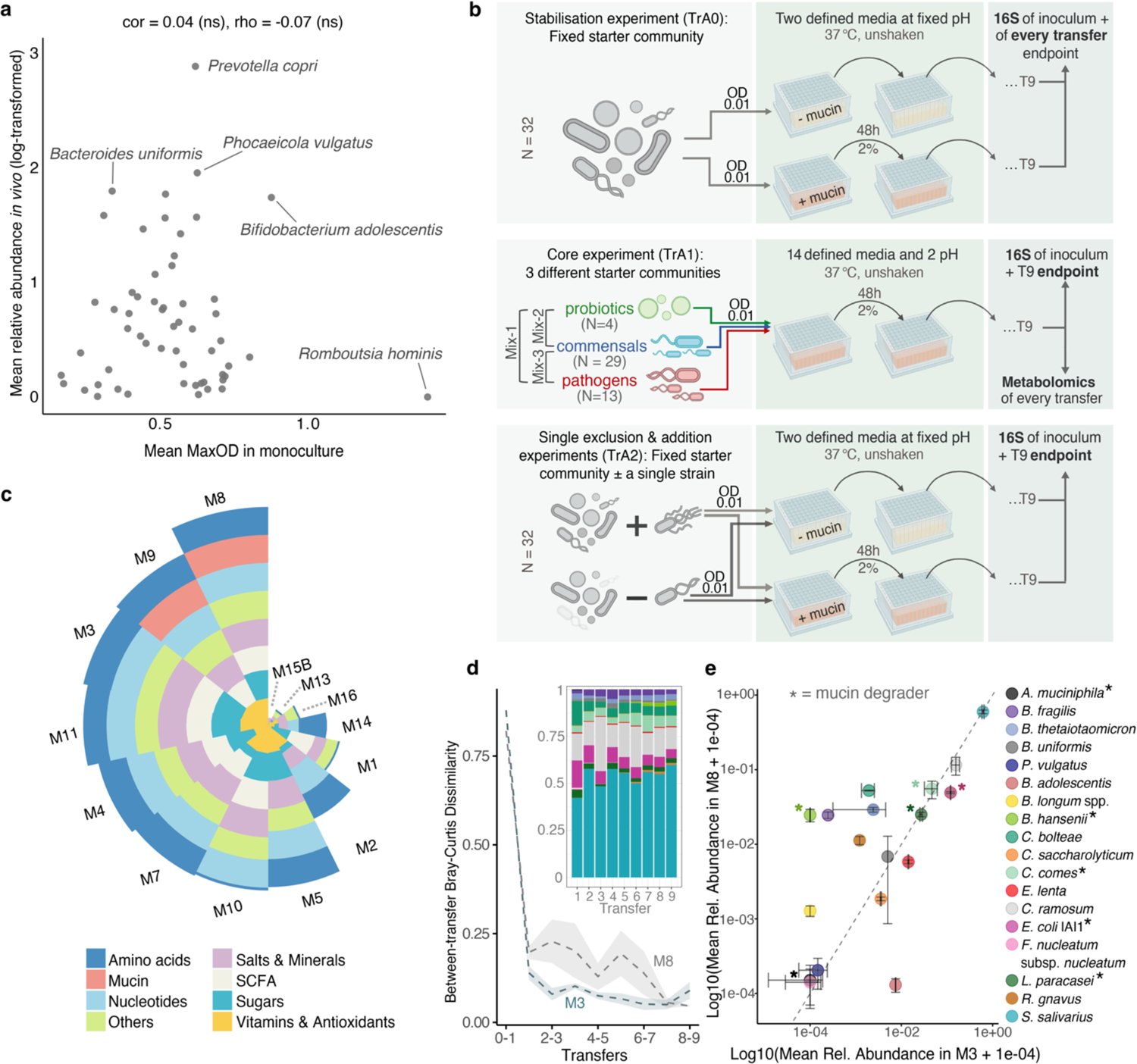
Assembly of synthetic gut bacterial communities to address the discrepancy between individual and community-level fitness. **a.** Relative abundances of gut bacteria *in vivo* do not correlate with their *in vitro* growth fitness in monoculture. *In vivo* relative abundances estimated from 5538 stool samples from healthy adults ^57^; *in vitro* fitness from ^10^. **b.** Overview of the serial transfer assembly (TrA) experiments. **c**. Overview of the defined media used in the study. The stacked bar-plot shows normalised medium components per nutrient class. If a medium contained the highest relative concentration of all components belonging to a nutrient class, its assigned value for that class was 1. **d.** Bray-Curtis dissimilarity between transfers showcases rapid stabilisation of the communities, with the mucin-enriched medium (M8) showing higher variability between transfers prior to stabilisation than the mucin-deprived medium (M3). The dashed line represents the mean trend, and the shaded area marks the standard deviation. Inset depicts the dynamics of the relative abundance changes for medium M8 (colour key as in **e**). **e.** Comparison of endpoint relative abundances between M3 and M8 media (TrA0 experiment). Asterixes highlight strains that utilise mucin in monoculture in accordance with ^10^. The dashed diagonal line marks equal relative abundance in M3 and M8. Only strains with nonzero relative abundance in either M3 or M8 are shown.

The resources essential for survival in a community must come either directly from the external supply (diet or host-supplied nutrients) or indirectly through the metabolic activity of the community members. While discerning between these two sources is difficult in *in vivo* settings where available nutrients reaching the community are hard to track, *in vitro* synthetic communities provide a tractable approach to delineate the contribution of the biotic supply. Previous work has shown that synthetic gut bacterial communities can be readily assembled through serial dilution and remain stable over multiple transfers^7–9^. However, these communities were assembled in undefined growth media obscuring the relation between specific nutrient components and community composition. To overcome this, we assembled gut bacterial communities using a set of 14 defined growth media with a gradient of nutrient richness. These media have previously been shown to support monoculture growth of circa 100 strains representative of healthy human microbiota ^10^. Since pH is a crucial factor for bacterial growth affecting many biochemical processes, including substrate utilisation, membrane transport, and ion homeostasis^11^, we assembled communities at pH 5.5 and 7, consistent with the pH range within the GI tract ^12^.

Biotic interactions such as resource competition ^13,14^, cross-feeding ^15–17^, and active antagonism ^18,19^ are determined by the membership of the community. Therefore, we also probed the effects of founding membership on community assembly using three inoculum compositions as well as systematic single strain exclusions and additions. Together, this comprehensive experimental design (132 distinct assemblies in total) allowed us to jointly investigate the role of abiotic (nutrients and pH) and biotic interactions on community composition.

## RESULTS

We started with a set of 63 well-characterised bacterial strains that are phylogenetically and metabolically representative of healthy human microbiota ^10^. Individually grown species were pooled together at approximately equal optical densities and the mixture was used as a community inoculum. Compositional dynamics were tracked using 16S amplicon sequencing and metabolic dynamics using untargeted mass-spectrometry of the co-culture supernatants.

The overall study covered three transfer assembly experiments (Fig. 1b, Suppl. fig. 1): i) TrA0: for testing the compositional stability over nine transfers in two defined media differing by mucin. ii) TrA1: the core experiment covering 84 combinations of 14 media, two pH, and three inoculum mixes; iii) TrA2: testing the impact of single species additions and exclusions (46 in total) from the inoculum mix for two defined media differing by mucin.

The three inoculum mixes used for the core experiment (TrA1) were: mix-1, consisting of commensals, pathogens, and probiotics; mix-2, containing commensals and additional probiotic strains; and mix-3, consisting of commensals and additional pathogenic strains (Fig. 1b, Suppl. fig. 1). These community mixes were assembled in 14 defined media, which comprised of several nutrient classes including sugars, amino acids, vitamins, salts, mucin, and short chain fatty acids (Fig. 1c), all of which are of relevance to microbiota diversity *in vivo* ^20–22^. Qualitatively, five media (M15B, M13, M16, M14, M1) were on the poorer side, and the rest (M2, M5, M10, M7, M4, M11, M3, M9, M8) on the richer side of the spectrum (Fig. 1c). Together, these media span a spectrum of nutrient richness within the constraints of the requirements of the least and most fastidious species.

### Communities rapidly assemble into stable compositional and metabolic state

To investigate stabilisation in defined media, we tracked community composition in media M3 and M8 – differing only by mucin – over nine transfers (experiment TrA0). The composition rapidly converged in both media stabilising by the fourth transfer (Fig. 1d). Presence of mucin in M8 enabled survival of mucin-metabolisers like *Blautia hansenii,* but also promoted several non-mucin-degraders (i.e., did not grow in monoculture with mucin as primary carbon source ^10^, e.g., *Bifidobacterium longum* spp.), indicating cross-feeding of mucin metabolism by-products (Fig. 1e, Suppl. fig. 2).

The stabilisation in four transfers we observed under defined media is consistent with those observed in complex, undefined growth media ^7–9,23^. However, stabilisation in taxonomic composition may not reflect functional stabilisation since composition does not capture potential shifts in metabolic activities of species and community. Since metabolic stability had not been addressed in previous studies (partly due to media complexity), we tracked community exometabolomes using mass-spectrometry of supernatants from our largest assembly experiment (TrA1, 84 communities over 9 time points) (Fig. 2a & Suppl. fig. 3). Metabolite profiles also stabilised from transfer 4 onwards (Fig. 2b & Suppl. fig. 3). These results attest the metabolic stability of the assemblies in the studied timescale enabling robust comparisons across different media and inoculation mixes.

**Figure 2.**
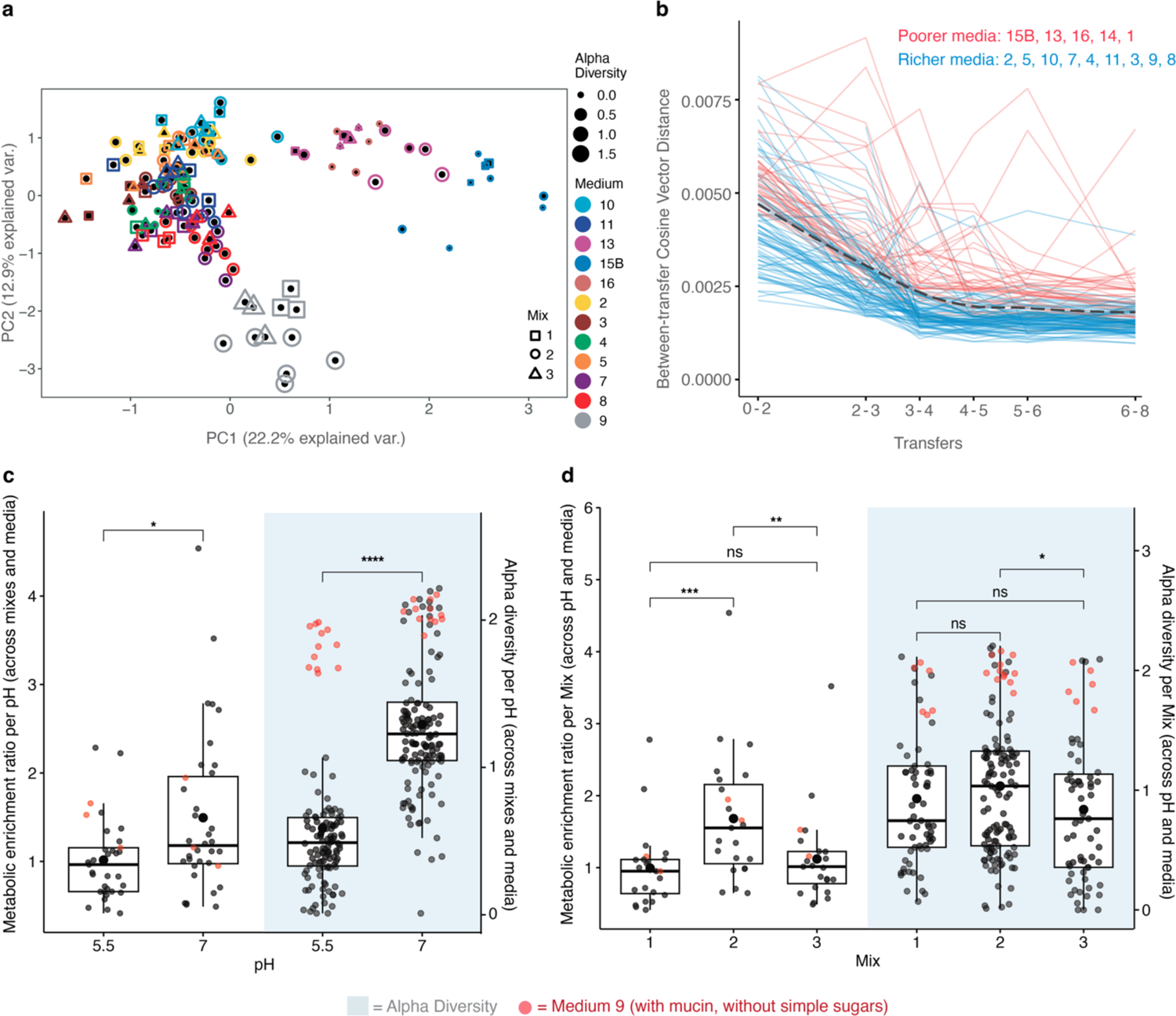
Exometabolome analysis highlights metabolic enrichment in community supernatants. **a**. Principal component analysis of untargeted metabolomics data (∼3500 ions) from pH 5.5 community supernatants highlights role of poor media (M15B, M16, M13) and mucin in the absence of simple sugars (M9) in shaping community metabolic landscapes. **b.** Rapid stabilisation of community supernatant metabolic profiles during assemblies (ph7, all medium and mixes). Shown are the cosine vector distances between metabolic profiles of the samples collected at consecutive transfers. The dashed line depicts the average trend (locally weighted smoothing). Timepoint 0 represents the empty media. **c.** Endpoint ratios of significantly enriched to depleted compounds and Shannon alpha diversity per pH (across all media and mix combinations). **d.** Endpoint ratios of significantly enriched to depleted compounds and Shannon alpha diversity per inoculation mix (across all pH and media combinations). For **c** and **d**, points in red mark medium M9 (mucin and no added sugars). Minimal media M15B and M16 are excluded as these predominantly harboured *E. coli* IAI1. Significance was determined using the Mann-Whitney U test with p-values adjusted using the Benjamini-Hochberg procedure. **** p < 0.0001; *** p < 0.001; ** p < 0.01; * p < 0.05; ‘ns’: not significant.

### Culture medium, pH, and inoculum mix jointly shape community assembly

Principal component analyses of relative abundance profiles of the 84 communities (experiment TrA1) show pH as a main driver of compositional variation with pH 5.5 generally leading to low-diversity communities (Fig. 2c, median final membership in pH 5.5 = 4, and in pH 7 = 10). At pH 5.5, communities were dominated either by *Lactobacillus paracasei* or *Lactobacillus plantarum*, two low-pH-tolerant species that competed with each other (Pearson’s R= −0.3, p=0.0005, Kendall’s tau = −0.2, p = 0.002) (Suppl. fig. 4). The two *Lactobacilli* drove abundance variability for most media except for M13, M15B and M9. M13 and M15B are the two most minimal media in our set, with M15B being specific for *Escherichia coli* spp. Indeed, *E. coli* spp. was the dominant species in these two minimal media, which lack several amino acids for which the two lactic acid bacteria are auxotrophic. The assemblies at pH 5.5 are consistent with the low diversity of the small intestine, which is also characterised by low pH and the survival of lactic acid bacteria therein ^24^. Growth medium M9, which has no added sugars but contains mucin, supported a more diverse and evenly distributed community including mucin-utilisers *Coprococcus comes* and *Clostridium perfringens* S107, as well as non-mucin utilisers *Clostridium bolteae, Bacteroides fragilis* spp.*, Clostridium ramosum*. Diversity promotion by mucin, even at low pH, suggests its metabolism helps counteracting the pH stress.

At pH 7, which is representative of the colonic environment where most gut bacteria reside, the variability in the relative abundance profiles is primarily driven by medium composition (Fig. 3a). The domination by *Escherichia coli* spp. in nutritionally poor M15B and M16 (defined minimal media for *E. coli* and *Veillonella parvula,* respectively) contribute to the separation, like in pH 5.5. While the community compositions are generally similar per medium irrespective of the inoculum mix, few exceptions are notable: richer media do not necessarily share similar abundance profiles (Fig. 3a); richer medium M9 clustered more closely with minimal media M13 and M1, rather than with fellow diversity-supporting, rich media such as M3 and M5 (Fig. 3a). Further, the communities arising from M7 inoculated with mix-2 shared a closer resemblance with communities in minimal medium M14 than with the richer medium’s mix-2 and mix-3 counterparts. Altogether, relative abundance profiles at pH 7 were jointly driven by medium and inoculum composition.

**Figure 3.**
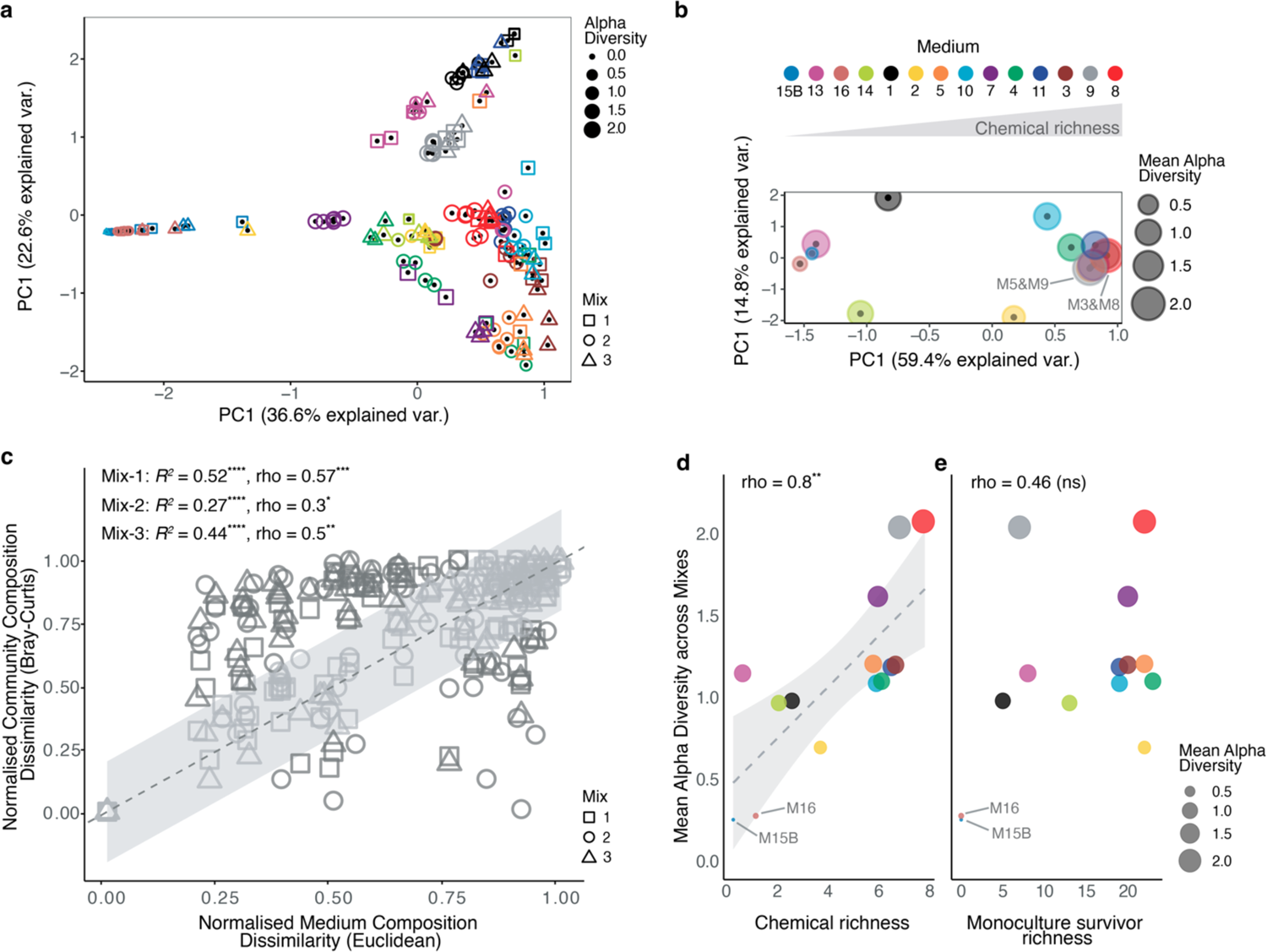
Medium composition only partly explains alpha- and beta-diversity of final assemblies. **a.** Principle component analysis of relative abundance data for pH 7 assemblies across different media and mixes highlight the contribution of medium richness to community diversity. **b.** Principle component analysis of the media components showcase how small differences in medium composition - even in richer media - have significant consequences for the alpha diversity of synthetic gut communities these defined media can support. Mean alpha diversity was estimated across the three inoculation mixes per media for pH 7. **c.** Medium (dis-) similarity (normalised Euclidean distance) is poorly correlated with the (dis-) similarity (normalised Bray-Curtis) between the supported communities. Data from final community composition (end of T9) for each of the three inoculation mixes used in the core experiment (TrA1) at pH 7. Grey-shaded area marks ± 0.2 from the diagonal with slope of 1 (dotted line). Significance of regression coefficients was determined using F-test. Significance for the Spearman rank correlations were estimated using Mantel test, 9999 permutations. **d.** Medium chemical richness corelated with the mean alpha diversity across mixes (at pH 7). Spearman’s rho = 0.798, p = 0.001; Pearson’s R = 0.762, p=0.002. **e.** Number of survivors supported by a medium in monoculture is a poor predictor of the alpha diversity. Spearman’s rho = 0.458, p = 0.099; Pearson’s R = 0.455, p = 0.102. **** p < 0.0001; *** p < 0.001; ** p < 0.01, * p < 0.05; ‘ns’: not significant.

### Biotic interactions underpin community diversity

To systematically assess the effect of medium composition on community membership, we used the sum of normalised components across classes as a metric of a medium’s chemical richness (Methods, Fig. 1c, 3b). The importance of any nutritional component to growth can vary across species due to their different metabolic needs. We therefore used an additional, biological, metric of medium complexity, viz. monoculture survivor richness, defined as the number of species a medium supports in monocultures (Methods). Surprisingly, while the chemical richness correlated strongly with the alpha diversity (Spearman’s rho of 0.80, p= 0.001, Fig. 3d), the biological metric – monoculture survivor richness – failed to do so (rho= 0.46, p= 0.1, Fig. 3e). Thus, a medium’s compatibility with individual species is insufficient in explaining whether those species would survive in the community and the overall co-culture diversity indicating a major role of inter-species interactions. Yet, a strong correlation between chemical richness and diversity hints at the metabolic nature of these interactions.

Amongst the individual chemical components, l-Cysteine, NAD, Hemin, Hematin, vitamins/antioxidants (Vitamin K, Lipoic acid, Biotin), and salts/minerals positively correlated with the alpha diversity supported by the medium (rho > 0.7, p < 0.01) (Suppl. table 2). Nevertheless, the difference in abiotic medium composition explains at most half of the variance in the relative abundance profiles (Fig. 3c), further emphasizing the contribution of metabolic interactions to final community composition. Yet, the strength of this correlation was inoculum mix-specific, with mix-2, which is enriched in probiotics, showing the lowest correlation (mix-1: rho = 0.57, p = 4e-04; mix-2: rho = 0.3, p = 0.03; mix-3: rho = 0.5, p = 0.002). This suggests that probiotics positively contribute to the effective nutritional environment for the community.

### Co-culture supernatants are metabolically enriched

Since all used media are chemically defined, the observed high diversity in low-monoculture-growth supporting media is likely due to additional, biotic, nutrient supply. To test this hypothesis, we assessed metabolite enrichment and depletion relative to the basal medium in co-culture supernatants using mass-spectrometry. We calculated the ratio between the number of ions with positive change to those with negative change (FDR corrected p<0.05 and |log2-FC| ≥ 1, Methods). For any co-culture exometabolome, a ratio of <1 means a net compound depletion while a ratio of >1 signifies net enrichment.

Across all co-cultures, pH 7 was found to be more conducive to metabolic enrichment than pH 5.5 (Mann-Whitney U test, p = 0.015, Fig. 2c), in line with the lower pH being energetically less favourable for most species. Probiotic-rich mix-2 featured stronger metabolic enrichment across pH regimes and media (Fig. 2d, Suppl. fig. 5). Mix-2 surpassed both mix-1 (Mann-Whitney U test, p = 0.0008) and mix-3 (Mann-Whitney U test, p = 0.007) in metabolic enrichment, while the difference between mix-1 and 3 was insignificant (Fig. 2d). The higher metabolic enrichment together with the higher alpha diversity in probiotic-rich mix-2 communities is consistent with our hypothesis that metabolic enrichment underlies diversity. In contrast, pathogens seem to offset the enrichment in mix-1 and 3 (Fig. 2d), consistent with the fastidious nature of these species ^25,26^.

To link species to the altered metabolic profiles, we constructed correlation networks between species relative abundances and metabolite changes. The metabolic landscape appears to be sculpted by a select number of species (Suppl. fig. 6). At pH 5.5, the primary players were, as expected from the abundance profiles, *E. coli* spp.*, L. paracasei*, and *L. plantarum*. At pH 7, which featured higher metabolic enrichment, many more (>10) species contributed to the network (i.e., having >5 significant correlations with ions). For example, *C. ramosum*, the species with most correlations (with ∼100 ions), is linked to several compounds that also positively correlated with other species. Together with the metabolic enrichment and diversity promotion, the observed species-ion correlations support cross-feeding as a driver of community assembly.

### Medium richness and communal support determine species’ success

To assess how species perform within a community relative to that expected from monoculture growth, we categorised species into two groups: beneficiaries and suppressed. Beneficiary species were those that performed better than expected (‘boosted’) in communities as well as the ‘emergent survivors’ – species that grew in co-culture despite failing to grow in monoculture in the same growth medium. Suppressed species are comprised of those that performed worse than expected (‘subdued’) as well as those that did not survive despite growing in monoculture (‘emergent extinction’) (Fig. 4a, Methods).

**Figure 4.**
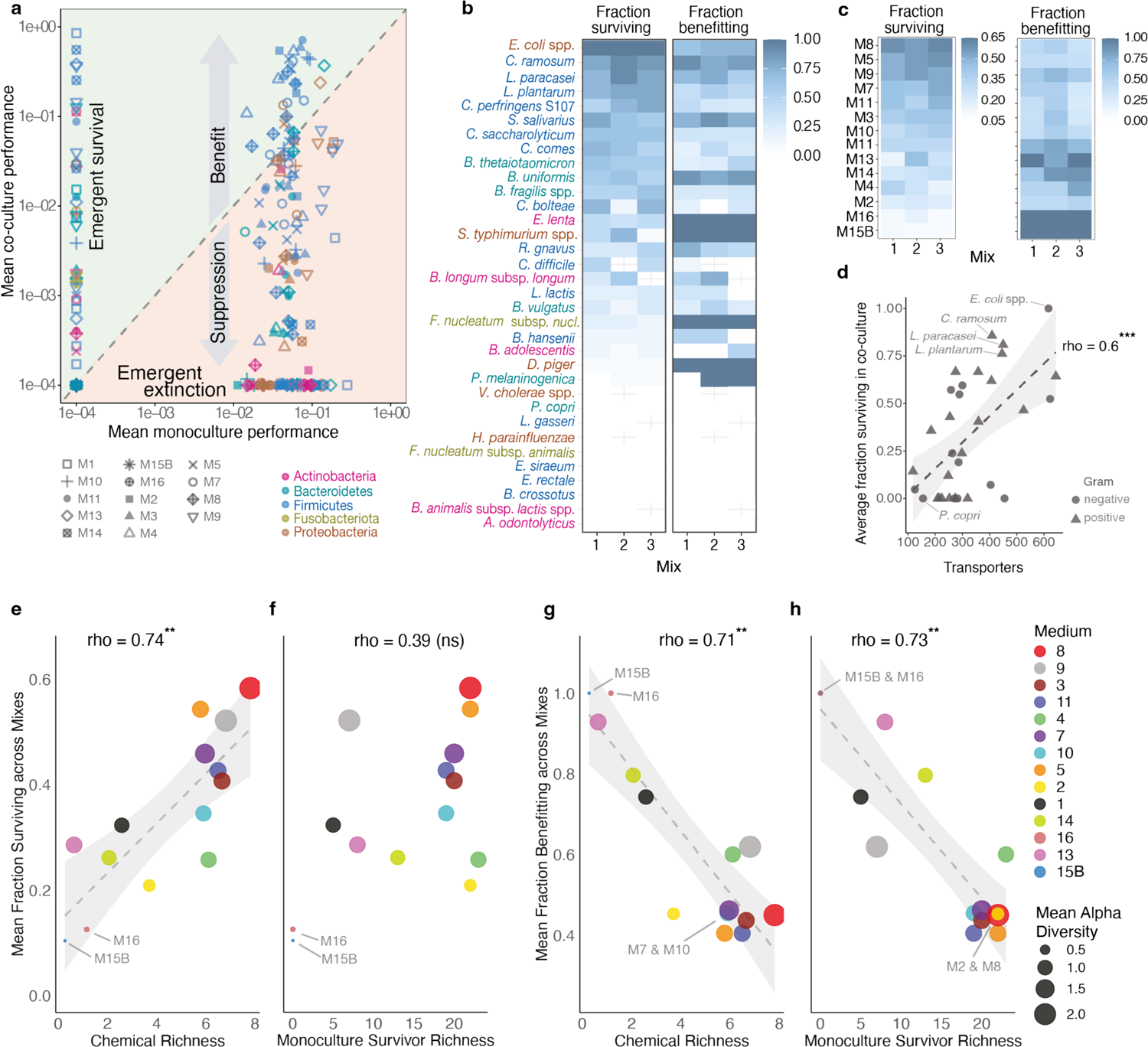
Emergence of beneficiaries and suppressed in community context. **a.** Species performance in monocultures (mean relative maxOD) does not correlate with their mean relative abundance in communities (inoculation mix-1 from the core experiment, pH 7). The diagonal marks the expected co-culture performance based on extrapolations from monoculture performance relative to that of other species present in the same inoculum in the respective medium tested. Both axes are in log-scale with all data points being adjusted with addition of a small number (1E-04) to account for 0 values (e.g. absent species) for visualization purpose. **b.** Heatmaps showing the fraction of species surviving, *F*_S_(’Emergent survivor’ + ‘Boosted’ + ‘Subdued’ + ‘Variable’) / (’No growth’ + ‘Emergent extinction’ + ‘Emergent survivor’ + ‘Boosted’ + ‘Subdued’ + ‘Variable’) and fraction of species benefitting, *F*_B_(‘Emergent survivor’ + ‘Boosted’) / (‘Emergent survivor’ + ‘Boosted’ + ‘Subdued’ + ‘Variable’), in co-culture. **c.** Heatmaps depicting the fraction surviving, *F*_S_, and fraction benefitting, *F*_B_, in co-culture for species across media. Empty cells mark taxa absent from the respective inoculum mix. **d.** Correlation of each strain’s predicted cytoplasmic membrane transport protein complements against its mean fraction survival and mean fraction benefitting across media and inoculation mixes. Fraction survival of species, averaged across media and inoculum mixes, correlated significantly with the number of predicted transporters (*F*_B_: Spearman’s rho = 0.56, p = 0.0006; *F*_S_: Spearman’s rho = 0.28, p = 0.10). Excluded from this figure (but not from correlations) is outlier *C. bolteae.* **e.** Correlation between chemical richness, the sum of all normalised media components (except for those in the ‘buffer’ category), and mean fraction species survival across mixes. **f.** Correlation between monoculture survivor richness of growth medium, with mean fraction survival across mixes. **g.** Correlation between chemical richness and fraction of mean fraction species benefitting across mixes. **h.** Correlation between monoculture survivor richness of growth media and mean fraction species benefitting across mixes. *** p < 0.001; ** p < 0.01; ‘ns’: not significant.

Across the 42 communities spanning three inoculum mixes and 14 media at pH7, 96% of the species were either beneficiaries or suppressed in at least one community. These included 153 cases (21%) of emergent survival, and 252 cases (35%) of emergent extinction. The chemically poorer media (M15B, M13, M16, M14, M1) were more conducive to emergent survivors, while rich(er) media (M2, M5, M10, M7, M4, M11, M3, M9, M8) featured more emergent extinction. On average, a poor medium had twice more emergent survivors than a rich(er) medium, while the latter had twice more emergent extinction (Suppl. table 1). The remarkable prevalence of emergent survival and extinction and their dependency on chemical richness highlights the role of metabolic interactions beyond resource competition.

A species’ propensity to survive or benefit in co-culture is not only dependent on growth medium, but also on the founding community composition (Fig. 4b). For instance, *L. paracasei*, a common probiotic strain, benefitted in co-cultures with additional probiotics (mix-2) more than in the other mixes (Fig. 4b, Suppl. table 3). Conversely, *C. perfringens* S107 displayed a greater survival in mix-3, which is low in probiotics (Fig. 4b). We note that the role of variation in inoculum load, i.e., difference in relative abundances of inoculated members, was not a substantive contributor as also observed previously ^23^ (Suppl. fig. 10). Overall, the differences regarding a species’ ability to survive and/or benefit across different community mixes in the same growth medium further underscores the role of communal support (or inhibition).

The prominent role of chemical complexity for overall species survival (Fig. 4c,e) considering metabolite enrichment (Fig. 2c,d) suggests that the ability to uptake diverse metabolites contributes to a species’ fate in a community. To test this, we correlated the predicted number of cross-membrane transport proteins, based on annotations by TransportDB and TransAAP ^27,28^, against a species’ performance in the community (Fig. 4d). Survival across media and mixes (*F*_!_) correlated positively with the number of predicted transporters (Spearman’s rho of 0.56, p = 0.0006) supporting cross-feeding in determining a species’ success. The correlation is maintained in a subset of species that feature survival in at least one condition (*F*_!_> 0; rho = 0.55, p = 0.005). However, *F*_!_did not correlate with genome size in this case (rho = 0.19, p = 0.38), suggesting that the link between transporters and co-culture survival is likely not determined by the genome size per se.

The propensity to support many species in co-culture appeared also to be dependent on the inoculum mix. For instance, across most media, mix-2 had a larger fraction of beneficiaries (except for M13, a minimal medium designed for *B. thetaiotaomicron*), attesting the key role of probiotics. Contrasting the overall survival and/or benefit between M9 and M8 is of particular interest since both are rich media with mucin, but M9 being free of (non-mucin-derived) simple sugars. While M8 was more supportive of survivors, M9 was more conducive of beneficiaries, especially for the mix containing no additional pathogens (mix-2), indicating a negating role of sugars against the diversity-boosting capacity of mucin (Fig. 2a, c, d).

While chemical richness positively correlated with the fraction of survivors (Fig. 4e,f), both chemical richness and monoculture survivor richness negatively correlated with the number of beneficiaries (Fig. 4g,h). The more species succeeded to grow as monoculture for any given medium, the more competition, and the fewer benefiters. This observation is not a direct extension of competitive exclusion, since a diverse community is retained after multiple transfers. The calculation of expected relative performance implies that any species that survive in monoculture should also survive in a co-culture; implicit in this calculation is (re)distribution of relative abundance space depending on how fellow members grow in respective monocultures. What we note here is that the observed degree of competition is disproportional between the rich and poor media. The contrast between poorer media – characterised by many beneficiaries – and rich media – characterised by many suppressed – shows how the trade-off between competitive and cooperative metabolism is modulated by the richness of the abiotic supply. The survival and benefit in a co-culture is thus jointly determined by the basal medium’s richness and how it is modified by the metabolic activities of community members.

### Species exclusions and additions attest biotic interactions

The composition of the inoculum and the presence of mucin seemed to drive exceptions to the trends otherwise explained by medium richness (Fig. 2a, Fig. 3d,e, Fig. 4c, e-h). To gain insights into how these two variables impact a species’ success, we investigated the effect of singly excluding or adding species to the inoculum in two media, one with and another without mucin (M8 and M3, respectively). Of all possible interactions between community members and added/excluded species (567 in M3, and 432 in M8), 42 (∼7%) significant (p< 0.05) responses were detected in M3, but twice more in M8 (81, ∼20%, Fig. 5a, Suppl. fig. 8). The majority (circa 40) of these additional responses in M8 were for non-mucin degrading species (*L. lactis, E. lenta, B. uniformis*, and *F. nucleatum* subsp. *nucleatum*), with only a smaller set (circa 10) recorded for mucin-degraders, viz. *A. muciniphila* and *B. hansenii*. Community-dependencies are thus largely driven by cross-feeding of mucin-degradation by-products rather than direct mucin utilization.

**Figure 5.**
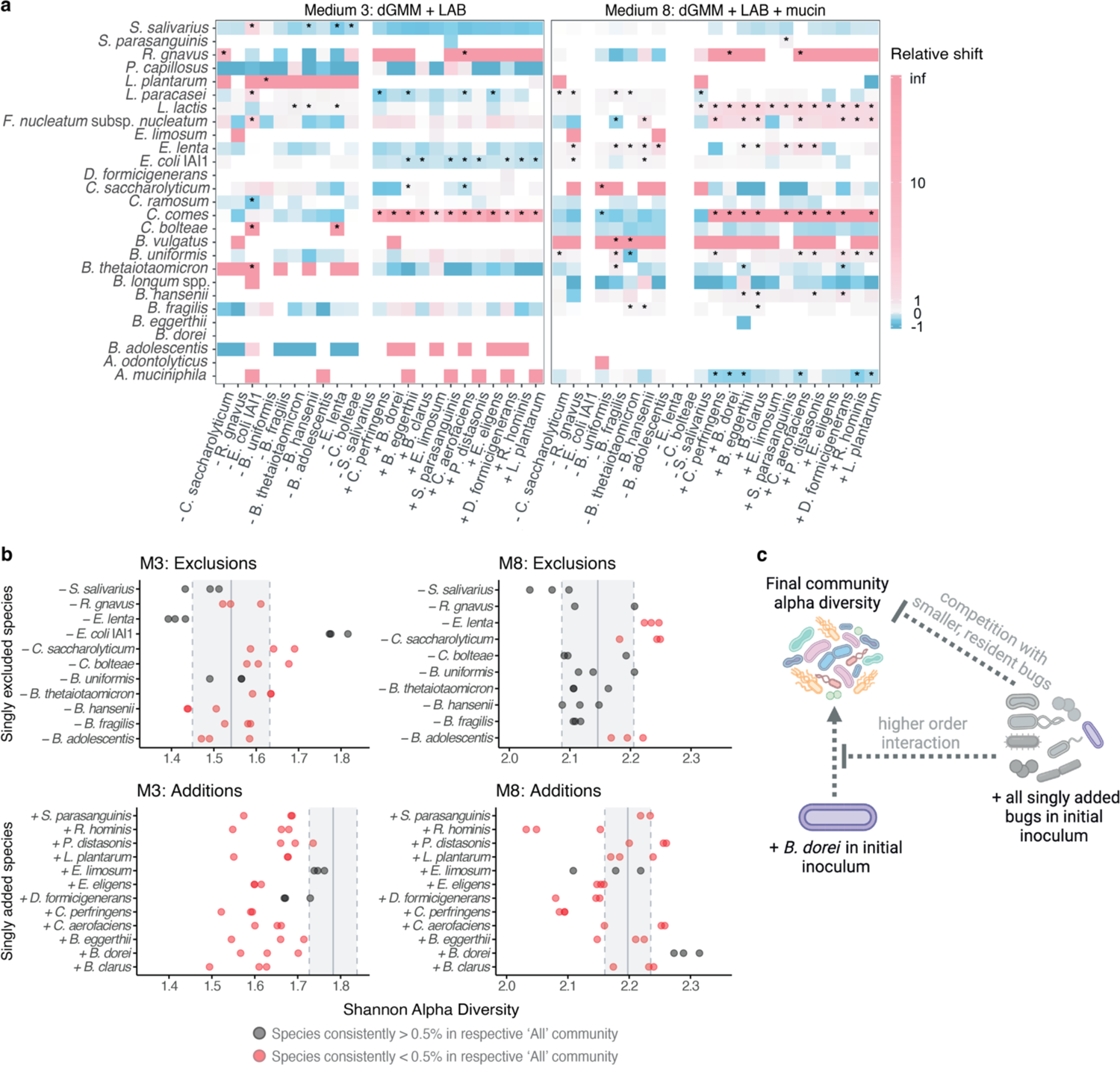
Species additions and exclusions show modulation of inter-species interactions by mucin. **a.** Impact of single exclusions, ‘-‘, and additions, ‘+’, on community members in M3 (left) and M8 (right) media. The colours denote shift in relative abundance compared to the baseline community including all species. Of all tested combinations of species and single exclusions or additions (567 in M3, 432 in M8), a total of 42 significant shifts (p< 0.05, marked with ‘*’) occurred as a response to single exclusions and/or additions in M3, and 81 in M8. **b.** Shannon alpha diversity indices for communities in M3 and M8 following single exclusion and additions. The vertical grey lines represent the mean (solid lines) and ± standard deviation (dashed lines) of the alpha diversity in the respective baseline community for exclusions (top panels) and additions (bottom panels). Single inclusion or exclusion of can have significant effect on the community diversity even if the added or excluded species in question has a low or near-zero abundance in the baseline community. **c.** Impact of *B. dorei* on community diversity as inferred from single addition/exclusion experiments. *B. dorei*’s alpha diversity-boosting effect in M8 was optimal when other singly added species were absent from the inoculum. Since *B. dorei* maintains a similar relative abundance in both the baseline (+’All’) as well its singly added scenario (∼2%), *B. dorei* itself does not appear to be suffering strongly from competition or other forms of ecological antagonism. Instead, when other singly added species are present in the inoculum, it may lead *B. dorei* to change its metabolism in a way that is less beneficial to low-abundant species. Alternatively, *B. dorei*’s shared resources, that would otherwise help some low-abundant strains gain more abundance, are consumed by any of the higher-abundant singly added strains in the ‘+ All’ baseline community and thus do not contribute to boosting diversity as they might when *B. dorei* alone is added to the inoculum.

The impact of inoculum exclusions and additions on community diversity was thus also modulated by mucin. For instance, the addition of *B. dorei* had a strong positive effect on alpha diversity in presence of mucin (M8) but not otherwise (Fig. 5b,c), despite *B. dorei* not being a mucin-degrader ^10^. Moreover, in M3, the exclusion of *E. lenta* resulted in reduced alpha diversity, while in M8, it increased the diversity despite *E. lenta* being a low-abundant member of the baseline community (Fig. 5b). This is consistent with the ecological importance of low-abundant strains ^29^, but emphasises its context-dependency.

A subset of species responded oppositely to single additions versus exclusions hinting at their general susceptibility to resource competition or benefit from common goods. *C. comes*, and to a lesser degree, *R. gnavus* and *F. nucleatum* subsp. *nucleatum,* generally benefitted from single additions and lost out in single exclusions. Yet, specific exceptions were also observed suggesting two-species interactions. For *F. nucleatum* subsp. *nucleatum*, a species linked to colon-cancer ^30,31^, exclusion of *B. hansenii* in M8 caused a positive shift while exclusion of *B. fragilis* caused a negative shift (Fig. 5a, Suppl. fig. 9). Similarly, while *S. salivarius* generally displayed negative shifts across single exclusions in M3, it benefitted from exclusion of *E. coli* IAI1. Most non-neutral effects of single additions and exclusions appear to be medium-specific as well. *E. coli* IAI1 had no significant response to any single addition in M8 but consistently lost in M3, while the opposite was observed for *L. lactis*. Together, the results from single strain additions and exclusions highlight the prevalence of inter-species interactions and their modulation by mucin.

### Logistic models partially capture dependency of co-culture survival on chemical richness

To what extent could the poor correlation between monoculture survivor richness and community diversity be due to the bottlenecking of slow growers? To answer this, we used ODE-based logistic growth models based on parameters estimated from monoculture growth kinetics and used these to simulate co-culture dynamics during the community assembly process (serial dilution). The simulations show that monoculture kinetics are informative of which strains will survive in community only for the very rich media, with circa 70% survivors correctly predicted for M8 and M5. An important exception is M9, a mucin-containing rich medium; only 10% of the survivors were correctly predicted. This further underscores the role of mucin in modulating biotic interactions, especially in the absence of sugars (Fig. 6a). As the models accounted for the bottlenecks during the assembly, the model predictions provide a better approximation for community survival than monoculture survivor richness (Fig. 6b, Fig. 4f). Yet, there remains a discrepancy that can only be justified via inter-species interactions.

**Figure 6.**
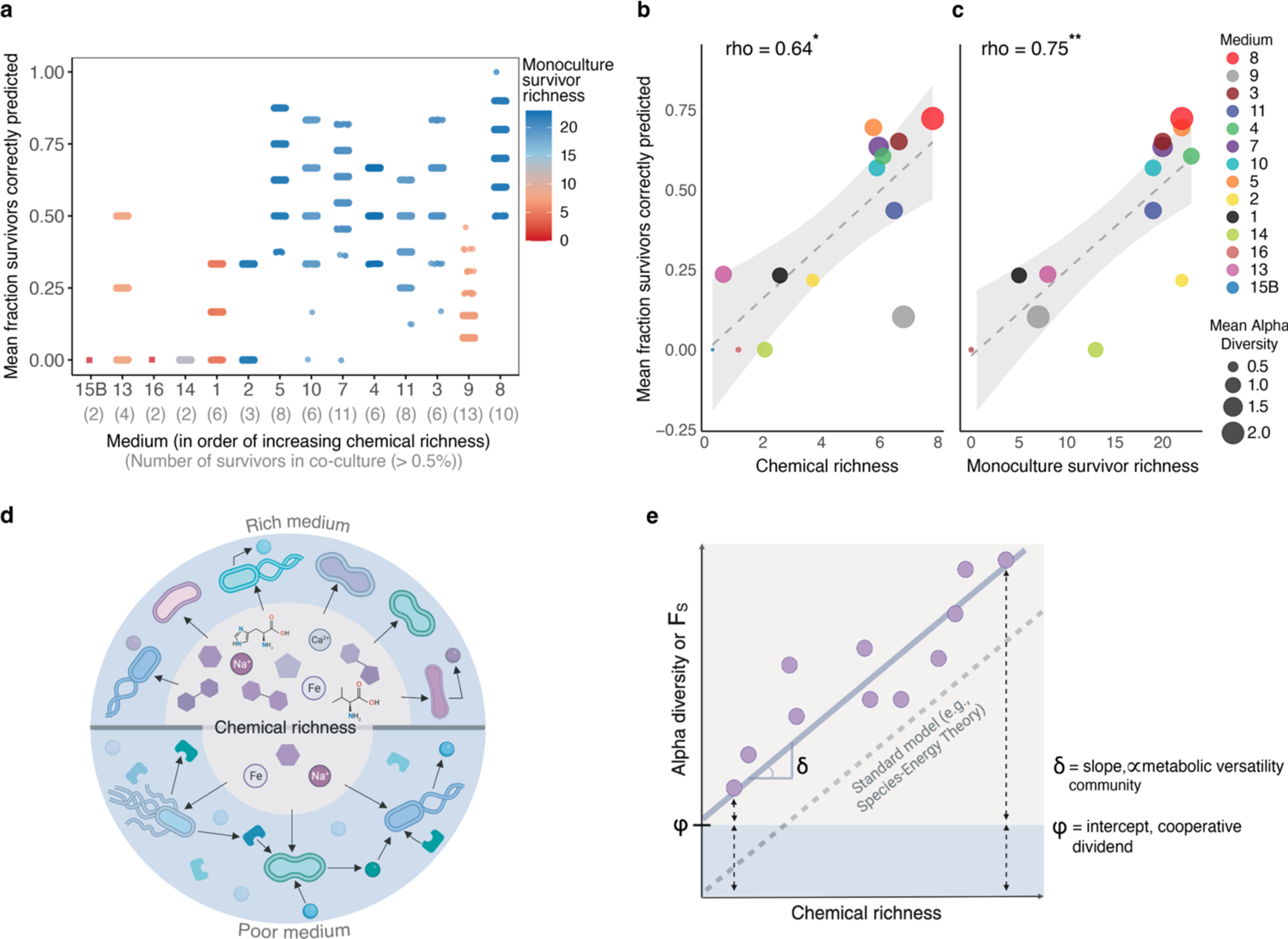
Cooperative dividend enables survival of species in poor media. (a) Fraction of co-culture survivors (i.e., >0.5% relative abundance) correctly predicted using an Ordinary Differential Equations (ODE) model informed by monoculture kinetics. We performed 1500 simulations per medium, with the outcome of each individual simulation corresponding to a single point in the plot. Neither M15B nor M16 supported any monoculture growth and hence were not simulated. **b, c.** The correlations between model performance (trained with monoculture kinetics) with two metrics of growth medium complexity: chemical richness (b) and monoculture survival richness (c). **d, e.** Emergent survival of many species in poorer media leads to a nonzero intercept (φ) for the linear relation between chemical richness and alpha diversity, or fraction survivors (*F*_!_). The slope (δ) of the relationship is proportional to community metabolic versatility/compatibility. Our data shows that the nonzero intercept emerges through metabolites secreted by the community members – hence termed cooperative dividend.

## DISCUSSION

While growth in a community inherently introduces competition for resources, it is becoming evident that cooperative forces also play a critical role ^32–34^. Through using defined culture media, our study provides a direct view into the role of competitive and cooperative forces in gut bacterial communities. A surprisingly large number of cases of both emergent survival (circa 20%) and emergent extinction (circa 35%) was observed in our study. While the shift towards extinction is consistent with resource limitation ^14,35,36^, we note the high fraction of emergent survival and the dependency on medium richness – emergent survival being more common in poorer media, while extinction in richer. Thus, we find ecological facilitation, especially cross-feeding, to be fundamental to the fate of most species, albeit modulated by the richness of the abiotic supply.

As the nutritional availability under the *in vivo* conditions is subject to fluctuating regimes driven by the host intake, both competitive and cooperative forces are likely to be important *in vivo*. The difficulty in mono-culturing a substantial fraction of microbiota members ^37,38^, even in highly rich media, further supports the role of cooperative interactions. We observe that the number of cross-membrane transporter genes encoded in a species’ genome positively correlated with its survival in co-cultures, providing a mechanistic support to the importance of cross-feeding.

The importance of mucin in creating cross-feeding opportunities was evident in our study as also observed previously ^39–41^. Our data additionally shows that this effect is contingent on the absence of simple sugars, which counter the mucin’s diversity boosting effect, offering an ecological explanation to *in vivo* studies showing negative impact of simple sugars on microbiota ^42,43^. In further support, we find that *B. dorei,* which positively contributed to alpha diversity in the presence of mucin but not in its absence, also correlates with alpha diversity across cohort studies (Suppl. table. 4).

The enrichment of supernatant metabolites and alpha diversity were both greater when inoculation contained probiotics. While the inclusion of pathogens had a negative effect on metabolite richness and community diversity, virtually all pathogens were undetectable in the final communities, emphasising the decisive role of transient community members as observed in other studies ^44,45^.

The results from the single strain addition and exclusion experiments points at the context dependence of the prevalence and complexity of higher-order interactions, which is still a topic of debate ^1,46,47^. The results also suggest strategies to counter potentially harmful strains such as colon-cancer associated *F. nucleatum* subsp. *nucleatum* ^30,31^: in the presence of mucin, *B. hansenii* emerged as a potential antagonist while *B. fragilis* as a potential helper (Suppl. fig. 9). Similarly, mucin-degrader *R. gnavus*, associated with Inflammatory Bowel Disease and Crohn’s disease ^48,49^, was strongly inhibited by the inclusion of *E. limosum* even though it benefitted from all other species. The role of emergent interactions is also evident in the discordance between monoculture and co-culture survival. The previous conjecture that the monoculture growth is predictive of community-scale performance ^46,50^ is thus not applicable to complex gut bacterial communities, neither *in vitro* when in poorer media (Fig. 4f, Fig. 6a) nor *in vivo* (Fig. 1a).

The biological metric of medium richness (number of species supported in monoculture) fared worse than the chemical richness (number of chemical components) in explaining emergent community phenotypes like alpha diversity. While the monoculture survivor richness captures individual species needs, it fails to account for the possibilities offered by community metabolism through metabolite secretion (Fig. 2b-d, ^33,51^). We conceptualise the latent niches embedded in the chemical richness as a cooperative dividend (Fig. 6d,e). Empirically, the dividend is observed as nonzero y-intercept of the linear regression between chemical and community richness (Fig. 3d). In richer media, more species can be supported by the abiotic supply, while the cooperative dividend becomes more apparent and consequential in poorer media. The non-zero y-intercept is also observed in other bacterial communities such as soil ^52^ supporting the universality of the cooperative dividend model. This extends the canonical ecological model of linearity between nutritional richness and community diversity, such as those within Tilman’s R* and Species-Energy Theory ^53–56^ by incorporating the role of inter-species interactions.

Overall, our study brings forward the fundamental role of emergent metabolic interactions in gut bacterial communities and their modulation by abiotic resources. Under nutritionally poor conditions, these interactions can manifest in communal metabolic dividend promoting emergent survival and diversity.

## METHODS

### Comparison of in vivo abundance with individual growth fitness

The average relative abundances of species were calculated from the 5.538 stool samples collected from healthy adult (age ≥ 18) included in curatedMetagenomicData (v3.8.0; ^57^).

### Gut bacterial strains and growth conditions

Bacteria were cultivated at 37°C under anaerobic conditions in a Vinyl Anaerobic Chamber (COY) inflated with a gas mix of approximately 15% carbon dioxide, 83% nitrogen and 2% hydrogen. Prior to the experiment, bacteria were pre-cultivated twice using one of the following media: modified Gifu Anaerobic Medium broth (mGAM, 05433, HyServe), Gut Microbiota Medium (GMM ^58^), Brain Heart Infusion broth (BHI, 53286, Sigma-Aldrich) supplemented with 2 mg/L NAD and 0.5mg/L Hemin (BHI++), MRS (69966-500G, Sigma-Aldrich) + 0.05% (w/v) Cysteine (MRS+), or mGAM supplemented with 10 mM taurine and 60 mM sodium formate (mGAM++). For long-term storage, a cryovial containing a freshly prepared bacterial culture plus 7% DMSO was tightly sealed and frozen at −80°C.

For the assembly of stable bacterial communities previously described defined and minimal media (MM) have been used: *B. thetaiotaomicron* MM ^59^, *C. perfringens* MM ^60^, *E. coli* MOPS MM1 and MM2 ^10,61^, *V. parvula* defined medium ^62^, Lactic Acid Bacteria medium (LAB ^63^), defined Gut Microbiota Medium (dGMM), dGMM+LAB and all recently published modified versions of the latter ^10^: dGMM+LAB containing only 10% (w/v) of amino acids, lower amounts of minerals and vitamins, monosaccharides or mucin as solely carbohydrate source, additional mucin and media excluding short chain fatty acids (SCFA) or aromatic amino acids. For complete media formulations, see Supplementary Materials in ^10^.

### Community assays

The defined starting communities used for all three experiments had most strains (i.e., 26) in common, while some strains were experiment specific (see Suppl. Table 3). The starting community used in the assembly stability experiment (TrA0) contained a total of 32 strains, while that of the core experiment (TrA1, mix-1) contained a total of 46 strains. The baseline for the single additions and exclusions experiment (TrA2) was identical to that of the assembly stability experiment, except for an additional 12 strains used for single additions. Considering the resolution of 16S amplicon sequencing, the different starting communities were composed of the following taxa: see Supplementary table 3 for overview.

### Sample collection and multiplexed 16S amplicon sequencing

To assemble a stable bacterial community, pre-cultures of individual strains were diluted in PBS to obtain an OD of 0.5 and mixed in a ratio of 1:1. Subsequently, the mixture was inoculated at an overall OD of 0.01 in 1 mL of the respective media in a 96 Polypropylene Deep Well plate (3959, Corning) sealed with a Breathe-Easy® sealing membrane (Z380059, Sigma-Aldrich). Every 48 hours, the culture was mixed, and 20 µL of culture were transferred to 1 mL of fresh media. To follow the community assembly process, at each time point of transfer, 100 µL of culture were transferred to a 96 MicroWell plate with Nunclon Delta Surface (163320, NUNC) sealed with a Breathe-Easy® sealing membrane (Z380059, Sigma-Aldrich). The pH was determined using non-bleeding MColorpHast pH indicator strips (Merck Millipore), and 500 µL of culture was pelleted and frozen at −80°C for subsequent DNA extraction and 16S amplicon sequencing.

To extract genomic DNA from 96 Polypropylene Deep Well plates (3959, Corning) containing the bacterial community samples, we adapted the GNOME DNA isolation Kit (MP Biomedicals) to be used with the Biomek® FXp Liquid Handling Automation Workstation (Beckman). Subsequently, purified DNA was obtained using ZR-96 DNA Clean & Concentrator™-5 (D4024, Zymo Research).

After the integrity of the DNA was verified by agarose gel electrophoresis, DNA concentration of the samples was determined using the Qubit dsDNA BR assay kit (Q32850, Invitrogen™) in combination with the Infinite® M1000 PRO plate reader (Tecan). The 16S V4 amplicons were generated using an Illumina-compatible 2-step PCR protocol: in a first PCR, the 16S V4 region was amplified with the primers F515/R806 ^64^, and then in a second PCR, barcode sequences were introduced using the NEXTflex 16S V4 Amplicon-Seq Kit (4201-05, Bioo Scientific).

After multiplexing equal volumes of PCR products from each sample, SPRIselect reagent kit (B23318, Beckman Coulter) was used for left-side size selection. Prior to Illumina sequencing the quality of the library was controlled using the 2100 BioAnalyzer (Agilent Technologies) and the DNA concentration was determined using the Qubit dsDNA HS assay kit.

Sequencing was performed using a 250 bp paired-end sequencing protocol on the Illumina MiSeq platform (Illumina, San Diego, USA) at the Genomics Core Facility, European Molecular Biology Laboratory, Heidelberg.

### Taxonomical assignments 16S amplicon sequencing

The raw data were filtered using fastp ^65^ (v 0.23.2) with default parameters, and samples with fewer than 2000 paired reads were excluded. Subsequently, the forward and reverse reads were merged using FLASH2 ^66^ (v2.2.00) with the following parameters: ‘-m 50 -M 150 -x 0.10’. The merged reads were then mapped to the reference sequence u sing Rbec ^67^ (v 1.1.4) with manual gene copy number correction.

### Relative Abundances Data Analysis

Relative abundance profiles of the different experiments were first explored using a Principal Component Analysis (PCA) using the ‘prcomp’ function of the ‘stats’ R package (v 4.2.2) and visualised using the ‘ggbiplot’ R package ^68^ (v 0.55). The Shannon alpha diversity index was calculated for all (endpoint) communities via: *−Σ^s^_i=1_ p_i_ • ln(p_i_)*, where *S* is the total number of species (with nonzero relative abundance), and *p*_i_ is the relative abundance of species *i*.

### The Core Experiment (TrA1, relevant for figure 2 d and figure 3)

We performed a redundancy analysis (RDA) to investigate the relationship between community composition and environmental variables ^69^. Species relative abundance data were treated as response variables, while variables pH, Medium, Mix and Replicate, were considered as explanatory variables. The RDA was conducted using the ‘rda’ function of the ‘vegan’ R package ^70,71^ (v 2.6.4). The significance of the model and the explanatory variables was assessed using permutation tests with 999 permutations. The variance explained by the model was partitioned into the individual contributions of each explanatory variable and their combined effects. Variance partitioning was then performed using the ‘varpart’ function from the ‘vegan’ R package, decomposing the total variance explained by the full model into fractions attributable to each explanatory variable and their overlaps.

To explore the role of biotic interference in the connection between the nutritional environment and community composition, we plotted medium dissimilarity against community dissimilarity. We calculated the Euclidean distance of normalised medium composition and the Bray-Curtis dissimilarity of final community compositions (end of T9 at pH 7) for each of the three inoculation mixes (i.e., we generated one matrix per mix, with each entry corresponding to a distance between two media; community composition of *M*^mix y^_i_ versus *M*^mix y^_j_). To calculate the Euclidean distance matrix, we used the ‘dist’ function of the ‘stats’ R package (v 4.2.2). To calculate the Bray-Curtis dissimilarity matrices, we calculated the mean abundances for each species in each combination of medium and inoculation mix and used the ‘vegdist’ function of the ‘vegan’ R package. To standardise the data within the matrices, we applied a normalisation function that scales the matrix elements by the maximum value in the matrix, effectively converting all entries to a range between 0 and 1. We calculated the R^2^ to quantify the relationship between the normalised medium composition distance (Euclidean) and the normalised community composition dissimilarity (Bray-Curtis) for each of the mixes and performed a t-test on the regression coefficients).

We correlated the chemical richness distance matrix with each inoculation mix’s community composition dissimilarity matrix using the Mantel test, performed using the ‘vegan’ R package. Specifically, the Mantel test was applied to assess the Spearman’s rank correlation between the Euclidean distance matrix of medium composition and the dissimilarity matrices of endpoint relative abundance profiles for each respective inoculation mix. To ensure the reliability of the results, the Mantel tests were run with 9999 permutations, allowing for the assessment of the significance of the observed correlations through random resampling of the data.

For any growth medium, chemical richness was calculated by taking the sum of all normalised media components (except for those in the ‘buffer’ category). Monoculture survivor richness was calculated by taking the total number of survivors in monoculture (i.e., mean final maxOD of > 0). To evaluate the association between these medium-specific metrics and the Shannon alpha diversity (averaged across all inoculation mixes), we computed the Spearman’s rho correlation coefficients.

To assess relative performance in monoculture versus co-culture, we calculated the mean relative maxOD for each species in each medium as follows: 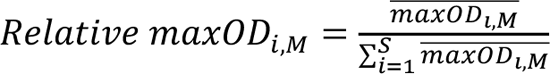, where 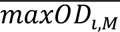 is the mean maximum optical density of species *i* in monoculture in medium *M* (where *i* ∈ (*i* … *S*)). This way, if two species performed equally well in monoculture, we expect that they would take up approximately equal proportions in co-culture when assuming neutral interaction effects and no 16S amplification bias.

To categorise a species’ discrepancy in relative growth performance in monoculture versus co-culture (for any specific medium), and to assess the consistency of this discrepancy across replicates, we developed a statistical classification method. Since the monocultures were not grown in low pH, we focused this analysis on relative abundance data extracted for pH 7. Since each inoculation mix has a unique initial community, we applied the status assignment for each inoculation mix separately (otherwise, a species that is absent from a certain mix might be falsely assigned an ‘emergent extinction’ status). The statuses, for any species in any combination of medium and inoculation mix, were defined as follows:

i. **No Growth**: when a species showed no growth in both monoculture and co-culture conditions.
ii. **Emergent survival**: when a species had a mean relative abundance greater than zero in co-culture while showing no growth in monoculture.
iii. **Emergent extinction**: Indicated when a species grew in monoculture but not in co-culture conditions.
iv. **Boosted**: when a species had nonzero mean (relative) abundance in both monoculture and co-culture, and when the upper bound of the 95% confidence interval of the mean relative abundance in co-culture exceeded the upper bound of the 95% confidence interval for monoculture.
v. **Subdued**: when a species had nonzero mean (relative) abundance in both monoculture and co-culture, and when the lower bound of the 95% confidence interval of the mean relative abundance in co-culture was less than the lower bound of the 95% confidence interval for monoculture.
vi. **Variable**: when a species had a nonzero mean (relative) abundance in both monoculture and co-culture but did not consistently meet the criteria for any of the above categories across replicates.

Following status assignment, we calculated the following fractions; either for species across media, or for media across species: fraction surviving 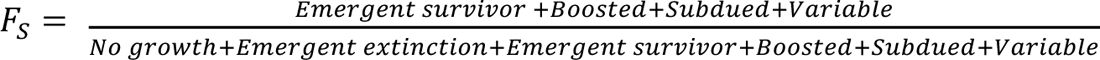, and fraction benefitting, 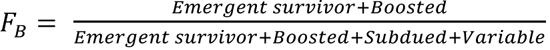. For a species’ fraction surviving, we account for the number of times this species consistently survived in co-culture across combinations of growth medium and inoculation mix. For fraction benefitting, we explore how much of survival is due to emergent survival or being boosted in co-culture. For a growth medium’s fraction surviving and benefitting, we account for the number of species consistently surviving or benefitting in co-culture when inoculated in this specific growth medium (across inoculation mixes).

### Correlation networks (relevant for Supplementary fig. 6)

Correlation networks between endpoint relative abundances of species and abundances of ions (of mix-2 samples from the core experiment) were constructed using respective Spearman correlation matrices, and visualised implementing the Fruchterman-Reingold force-directed algorithm algorithm of the ‘igraph’ R package ^72^ (v 1.4.2). Mix-2 was used due to the larger replicate space for end-point data (i.e., relative abundances and untargeted metabolomics of both endpoints; T8 and T9). Both endpoints were considered since inter-transfer variation was considered minimal (see Suppl. fig. 11).

### Single Additions and Exclusions Experiment (TrA2, relevant for figure 5)

The singly excluded species were generally species with high relative abundance in the community as observed for the assembly stability experiment for either M3 or M8 (Fig. 1 f). The singly added species were species that were absent from the baseline communities, and had high growth in monoculture (see Supplementary table 5 for monoculture growth, also included in ^10^).

Since the initial community composition for any single exclusion experiment and the baseline community differed by 1, our null expectation was that any species would benefit (in their relative abundance) from having this one species not present in the inoculum, provided that the singly excluded species occupied >0 relative abundance in the baseline community. To account for this ‘gap’, we calculated ‘discounted abundances’ to distill if a species responded particularly disproportionately to any single exclusion. Discounted abundances were calculated as follows:

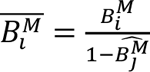, for *j = the singly excluded species*, where 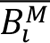 is the discounted abundance of species *i* in Medium *M*, *B*^M^_i_ is the relative abundance of species *i* in Medium *M*, and 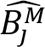 is the relative abundance of experimental species *j* in the baseline community (‘-All’) in medium *M*.

Since the initial community composition for any single addition experiment and the baseline community differed by *E* − 1 (the total number of singly added species minus the singly added species in question), we calculated the discounted abundance in single addition experiments as follows:

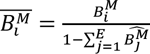, for *j* ≠ the singly added species, where *B^M^_i_* = the the discounted abundance of species *i* in Medium *M*, *B*^M^_i_ is the relative abundance of species *i* in Medium *M*, and *B*^M^_j_, is the relative abundance of experimental species *j* ∈ (1 … *E*) in the baseline community (‘+All’) in medium *M*.

From these discounted abundances, we calculated shifts induced by these individual experiments: *Shift = B^M^_i_ − B^M^_j_*, and 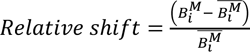. The latter was calculated to emphasise effects on species with a low to mid-tier abundance in the respective baseline community. In addition, we performed a t-test to assess whether measured relative abundances were significantly different from predicted discounted abundances for any combination of species, experiment, and medium. Shifts were then visualized using heatmaps.

Any species-consistent blanks within the heatmap can be attributed to species that failed to gain a non-zero relative abundance in either the respective baseline community or any single addition/exclusion communities in the respective medium. In single exclusion experiments, combinations in which a species was itself excluded were blanked. Similarly, for single additions, any single addition that was not that of the added species itself were blanked (as these would inherently lead to significant changes from the respective baseline communities). Unsuccessful single exclusion experiments (i.e., the excluded species itself was present in ≥2/3 replicates) were also disregarded. See Supplementary table 6 for overview.

### Metabolomics

Metabolomics analysis was performed as described previously ^73,74^. Briefly, samples were analysed on a LC/MS platform consisting of a Thermo Scientific Ultimate 3000 liquid chromatography system with autosampler temperature set to 10° C coupled to a Thermo Scientific Q-Exactive Plus Fourier transform mass spectrometer equipped with a heated electrospray ion source and operated in negative ionization mode. The isocratic flow rate was 150 μL/min of mobile phase consisting of 60:40% (v/v) isopropanol:water buffered with 1 mM ammonium fluoride at pH 9 and containing 10 nM taurocholic acid and 20 nM homotaurine as lock masses. Mass spectra were recorded in profile mode from 50 to 1,000 m/z with the following instrument settings: sheath gas, 35 a.u.; aux gas, 10 a.u.; aux gas heater, 200° C; sweep gas, 1 a.u.; spray voltage, −3 kV; capillary temperature, 250° C; S-lens RF level, 50 a.u; resolution, 70k @ 200 m/z; AGC target, 3×10E6 ions, max. inject time, 120 ms; acquisition duration, 60s. Spectral data processing was performed using an automated pipeline in R. Ions detected in less than 75% of samples were removed, and 3480 ions remained for further analysis. Ion intensity drift, associated with degrading instrument performance during data acquisition which occurred in randomized sample order, was corrected by a median filtering-based approach. Detected ions were tentatively annotated as metabolites based on matching accurate mass within a tolerance of 5 mDa using the Human Metabolome database as reference ^75^, assuming [M-H]^-^ and [M-2H]^2-^ as ion species as well as at most two ^12^C to ^13^C exchanges. Of note, the resulting annotations are only tentative and can lead to ambiguous metabolite assignments; for instance, isomers cannot be distinguished by this approach.

### Chemical Composition Normalisation (relevant for figure 1e)

Normalisation followed: 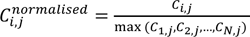, where *C^normalised^_i,j_* represents the normalised concentration of component *j* in medium *i*, *C*_i,j_ is the original concentration of component *j* in medium *i*, and max (*C*_1,j_, *C*_2,j_, …, *C*_N,j_) represents the maximum concentration of component *j* across all media: {*M*_1_, *M*_2_, …, *M*_N_}.

Normalised medium components were then summed per class (i.e., amino acids, nucleotides, salts and minerals, etc.). With mucin being one of the few complex organic molecules that is added to a subset of defined media (M9 and M8), classifying only these two media as ‘semi-defined’, mucin was taken as a separate medium component class. The normalised medium components summed per class were in turn normalised to arrive at a neutral measure of chemical richness (i.e., no bias in richness towards classes with more individual components). Hence, across all 8 chemical classes (Fig. 1e), the maximum chemical richness to be assigned is 8 (i.e., the medium would need to have the highest concentration across all of the components in each class).

### Statistical Analyses Untargeted Metabolomics

### Principal component analyses (relevant for figure 2a)

Principal Component Analyses (PCAs) were run on normalised untargeted metabolomics of final transfers (transfer 9 for Mix-1 and 3, transfer 8 and 9 for Mix-2) using the ‘prcomp’ function of the ‘stats’ R package (v 4.2.2).

### Cosine vector distances (relevant for Figure 2b)

We calculated the cosine vector distances of untargeted metabolomics measured at pH 7, across all transfers and mixes (from which supernatant samples were taken). Cosine vector distances were calculated as follows:

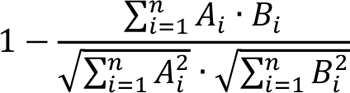

Where *A*_i_ and *B*_i_ are the components of vectors *A* and *B*, respectively, for *i* = 1, 2, …, *n*, and where *n* is the number of components (dimensionality) of the vectors. The components correspond to distinct ions measured (a total of 3480), and vectors *A* and *B* correspond to sequential transfers compared (i.e., transfer 3 versus transfer 4) of a particular combination of Mix, Medium, pH and Replicate. We accounted for *t*_0_ by taking the metabolomics data of medium controls (i.e., without inoculation of any species) measured for each combination of Medium, pH and Replicate (and use this same vector of values as *t*_0_ for each Mix, Medium, pH and Replicate combination).

### Enrichment versus depletion analysis (relevant for Figure 2d)

We calculated log2 fold changes (Log2FCs) of metabolites at final timepoints (Transfer 9) relative to media controls (Transfer 0) for all conditions (i.e., combinations of pH, Medium and Inoculum mix) using the ‘edgeR’ R package ^76^ (v 3.40.2). Significant Log2FC values (i.e., FDR corrected p < 0.05) were then used for further analysis, including PCA.

Per combination of inoculum Mix, pH and Medium, we summed all compounds that were significantly depleted or enriched, and statistical comparisons between pH regimes (across all Media and Mixes) and between Mixes (across all Media and pH) were performed using a Mann-Whitney U test (with Benjamini-Hochberg method for adjusting the p-values).

### Putative metabolite annotations (relevant for supplementary figure 12)

Not every significantly enriched or depleted ion (for any combination of Medium, pH and Mix) was annotated following the annotation method described under Metabolomics, whereas some ions have multiple annotations at the chosen chemical taxonomical resolution: compound class. If a significantly depleted or enriched ion was annotated across *N* classes, its weight within that class would be divided by *N* so that *1/N* would be attributed to the total count of compounds significantly enriched or depleted in compound class *y*. Then, for each compound class, we subtracted the total score of depleted compounds from the total score of enriched compounds, to arrive at an estimation of enrichment versus depletion for any compound class for each unique combination of Medium, pH and Mix (i.e., if >0, more compounds tended to be enriched for that specific compound class).

This analysis was limited to providing insights into relative enrichment versus depletion per compound class for ions that were annotated, and thus may not be representative of the true distribution of enrichment versus depletion per compound class. To check the accuracy and precision of our metabolomics annotations and to validate the robustness of our approach in distinguishing between different classes of compounds, we conducted a comprehensive comparison of the metabolic profiles of various growth media (see further Methods in supplementary figure 12).

### Modelling

We simulated the logistic growth curves in a serial dilution regime for all 34 taxonomically distinct strains of Mix-1 (core experiment), with parameter inference from monoculture kinetics (averaged across strains belonging to taxonomically distinct species groups).

A system of 34 coupled ordinary differential equations (ODEs) was formulated as follows: 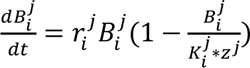, where *r*_i__j_ is the maximum growth rate for species *i* in medium *j*, pulled from normal distribution of monoculture kinetics (*r*^j^_i_∼N(µ(*r*^j^_i_), σ(*r*^j^_i_)), and where *K*^j^_i_ is the maximum optical density (MaxOD, i.e., species-specific carrying capacity) for species *i* in medium *j*, pulled from normal distribution of monoculture kinetics (*r*^j^_i_∼N(µ(*K*^j^_i_), σ(*K*^j^_i_)). We implemented a co-culture correction factor, *z*^j^ = c^j^, where *c*^j^_i_ is the sum of maxODs observed in monoculture for community members followed in medium *j*, and *m*^j^ is the maximum maxOD observed across monocultures for medium *j*. This co-culture correction, which is medium-specific, forms a null-hypothesis of proportional distribution of abundance in co-culture, and thereby implicitly accounts for a degree of universal ‘abiotic’ competition imposed by limited ‘space’.

The model was integrated with initial conditions inferred as follows: per simulation, we have drawn the relative abundance of species *i* from a normal distribution defined by its variability observed across the inoculum replicates: (*B*_#%_ ∼N(µ(*B*_#%_ ∗ 0.01), σ(*B*_#%_ ∗ 0.01)^R^)). The multiplication by 0.01 transformed this normal distribution from relative abundance to OD units (since the OD of initial mixture was ∼ 0.01). Further, model integration accounted for the experimental design, defined by a dilution regime of 9 transfers of 48 hours with 2% transferred volume.

To account for the variability in parametrization in full combinatorial space (i.e., each species’ growth kinetics and initial densities were drawn from respective normal distributions per simulation), 1500 simulations were run per growth medium. The simulation output was analysed as follows:

1. The endpoint (432h) values were turned into relative abundances since simulated *B*\* is in OD units;
2. We then counted, per simulation, how many simulated survivors were correctly predicted to be survivors (i.e., had >0.5% abundance both in the simulation endpoint and in at least 2/3 replicates in the real relative abundance table belonging to endpoint data for medium *j*);
3. We finally calculated the fraction of survivors correctly predicted by dividing the correct survivor count by the total number of ‘real’ survivors in the relative abundance table for medium *j*

## Data and code availability

Relative Abundance data is uploaded to ENI (https://www.ebi.ac.uk/ena/) under this accession number PRJEB71340. Metabolomics data is submitted to MetaboLights (https://www.ebi.ac.uk/metabolights/) under the following accession number MTBLS9413. All code used for data analysis is available at GitHub (https://github.com/NaomiIrisvdBerg/Emergent_survival_and_extinction_gut_bacteria).

## Author contributions

M.T., S.A., A.T. and K.R.P. designed the study. M.T. performed microbiology experiments, DNA extraction and 16S rRNA library preparation. S.B., Anja T., M.K., and A.R.B contributed to microbiology and molecular biology experiments. R.G. and Y.K. analyzed the sequencing data. S.A. and L.K. helped with data analysis. D.S. performed metabolomics analysis. N.I.B. analyzed the data, performed modelling, and prepared figures. K.R.P., P.B., and A.T. supervised the experimental work. K.R.P. supervised data analysis and coordinated the overall study. N.I.B., M.T. and K.R.P. wrote the manuscript. All authors read and commented on the manuscript.

## Acknowledgements

Sequencing was performed at Genecore, EMBL. This project has received funding from the European Union’s Horizon 2020 research and innovation program under grant agreement no. 686070. M.T. was supported by the EMBL interdisciplinary postdoctoral program.

## 8. SUPPLEMENTARY FIGURES

**Supplementary figure 1:**
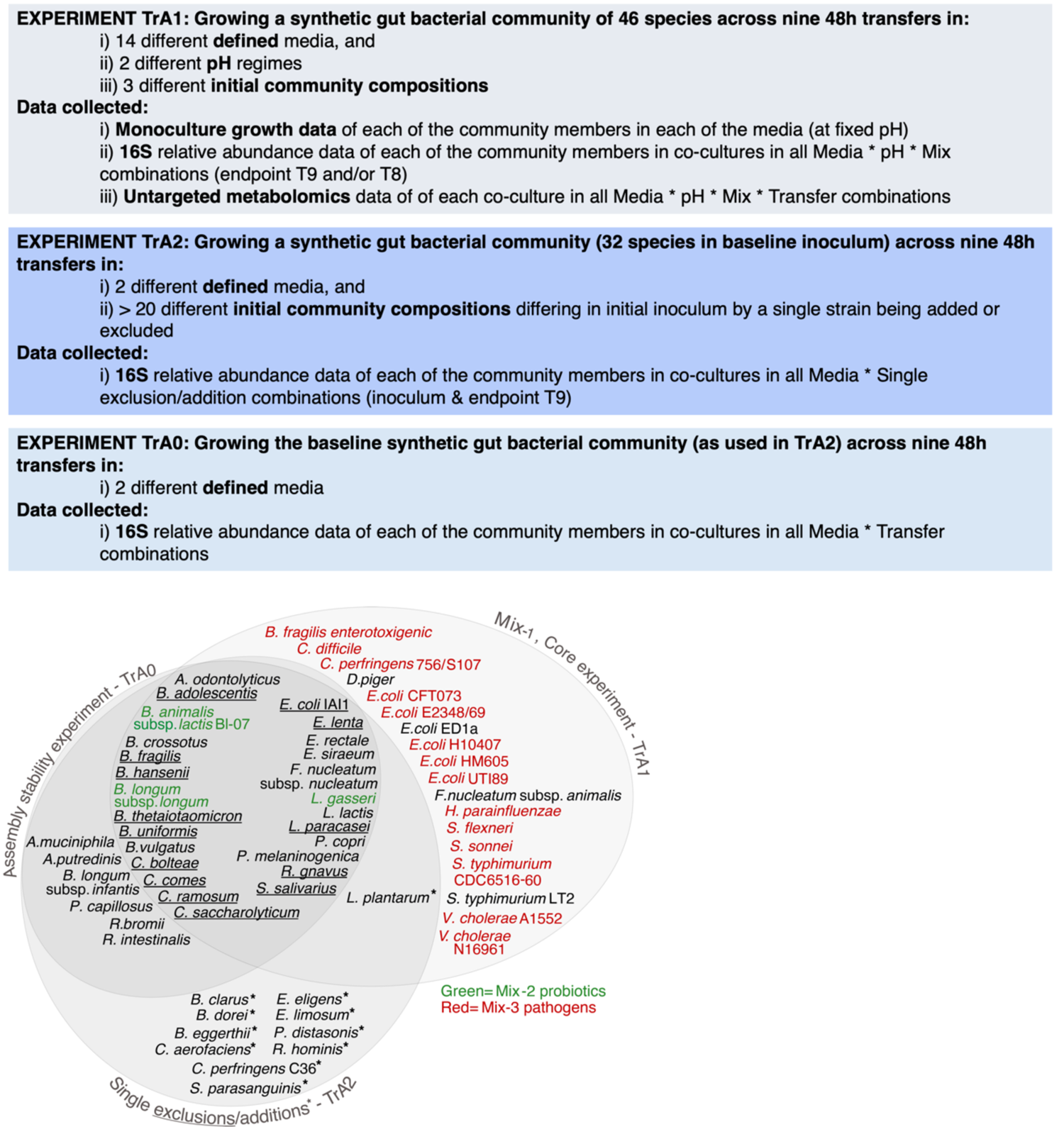
Overview of experimental design and generated data for all three analysed experiments. The Venn diagram contains an overview of the inoculum mixes used across the three experiments. Species underlined are those that are singly excluded in the TrA2 experiment, while those with an asterisk (*) highlight species that are singly added. See Supplementary Table 3 for a complete overview of inoculum compositions of each experiment.

**Supplementary figure 2.**
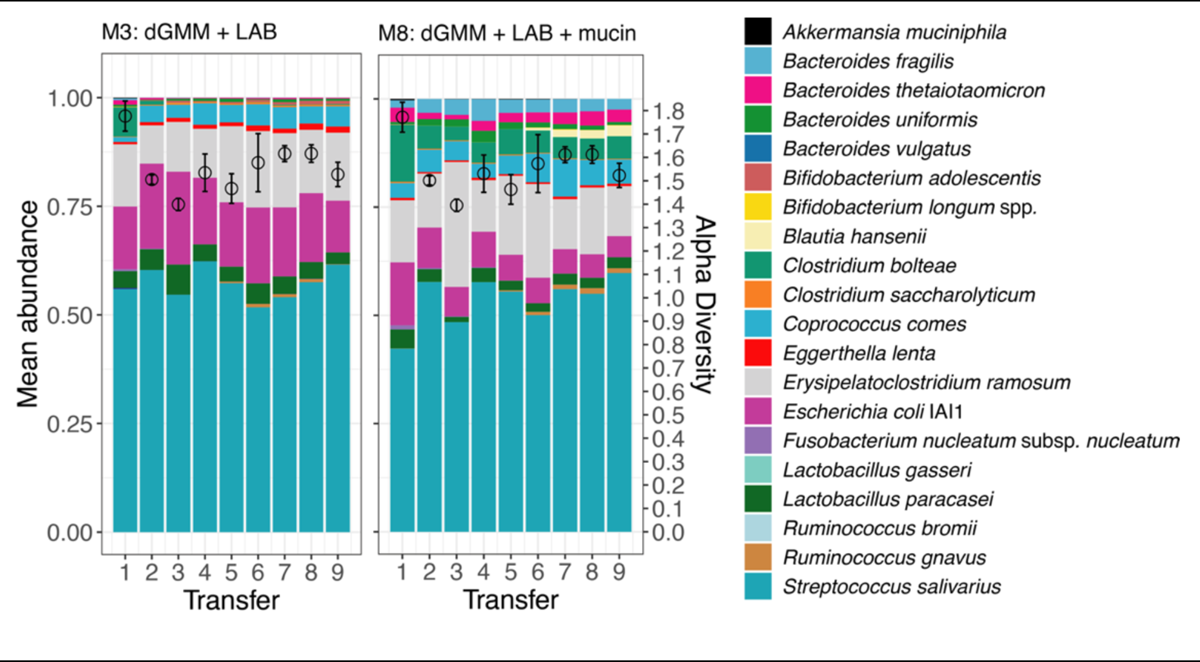
Relative abundances of bacterial strains in Medium 3 (M3) and Medium 8 (M8) across transfers.

**Supplementary figure 3.**
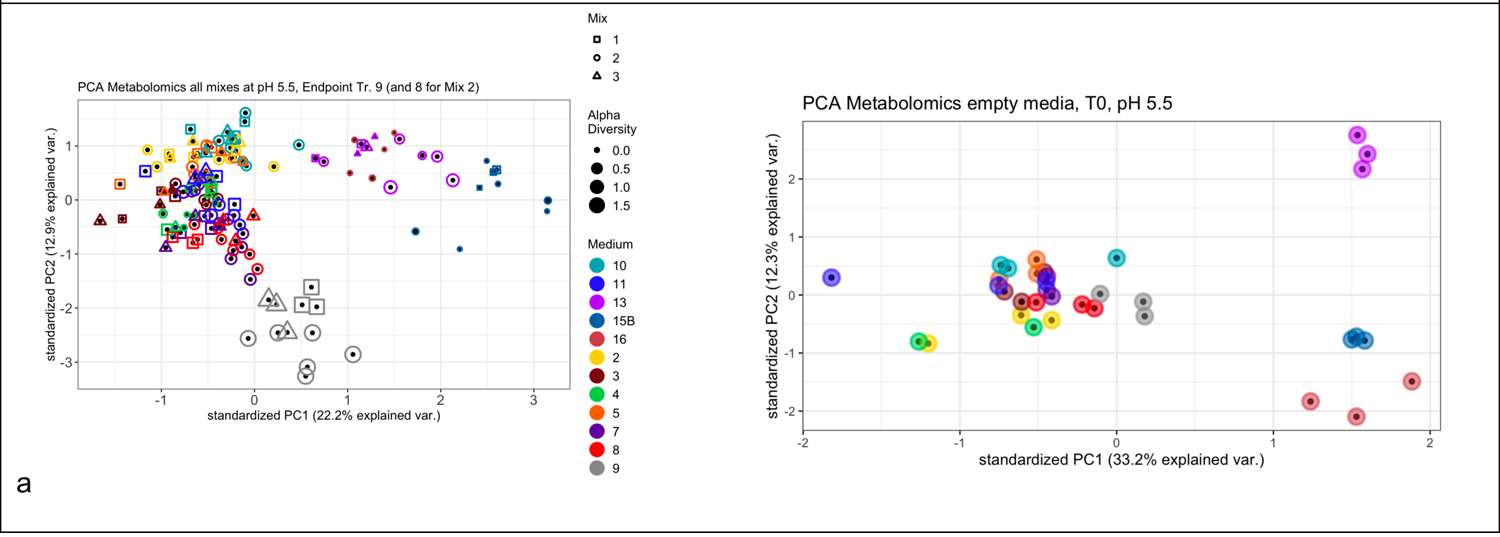
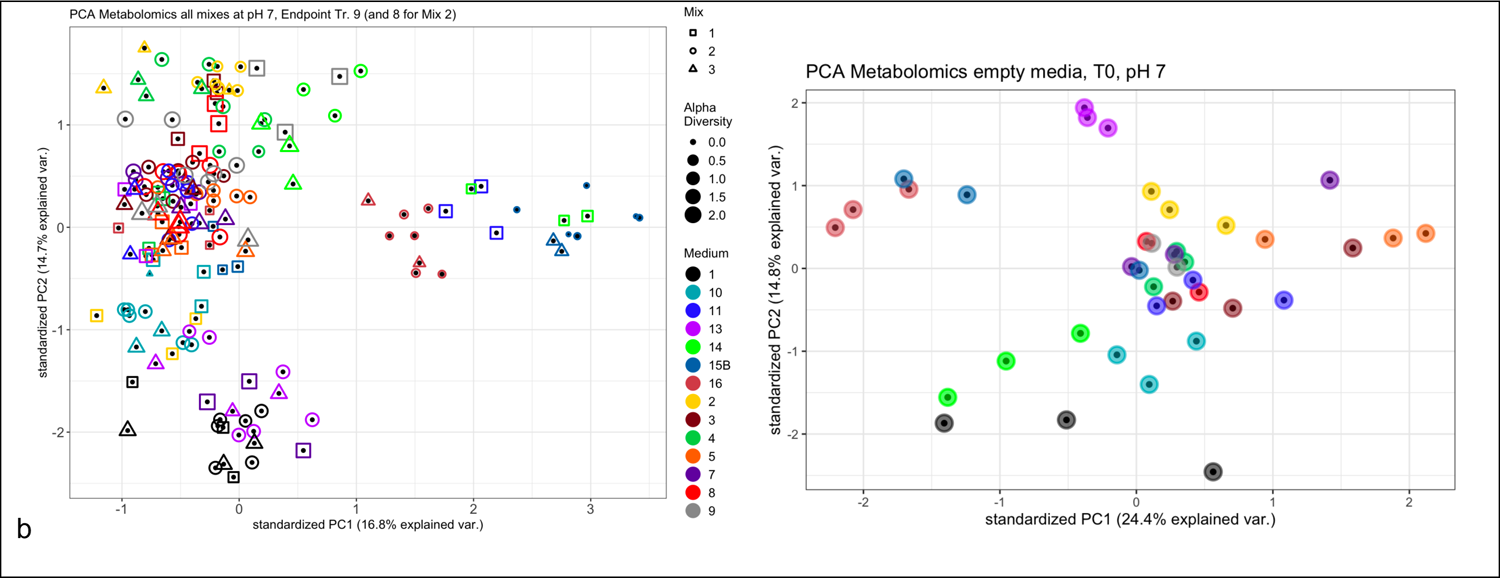
PCAs of normalised untargeted metabolomics data in pH 5.5 (a) and pH 7 (b). In accordance with 16S data, pH is the main driver of variability across all (∼3500) measured ions and is hence separated out. The variability in both PC1 of empty media versus endpoint co-culture supernatants is driven by poorer media M15B, M16, followed by M14, M13 and M1. This suggests that PC1 in untargeted metabolomics of endpoint supernatants is driven by medium formulation. For the T0 PCAs, we excluded the following outliers; in pH 5.5: Medium 4, replicate 3 and in pH 7: Medium 8, replicate 2.

**Supplementary figure 4.**
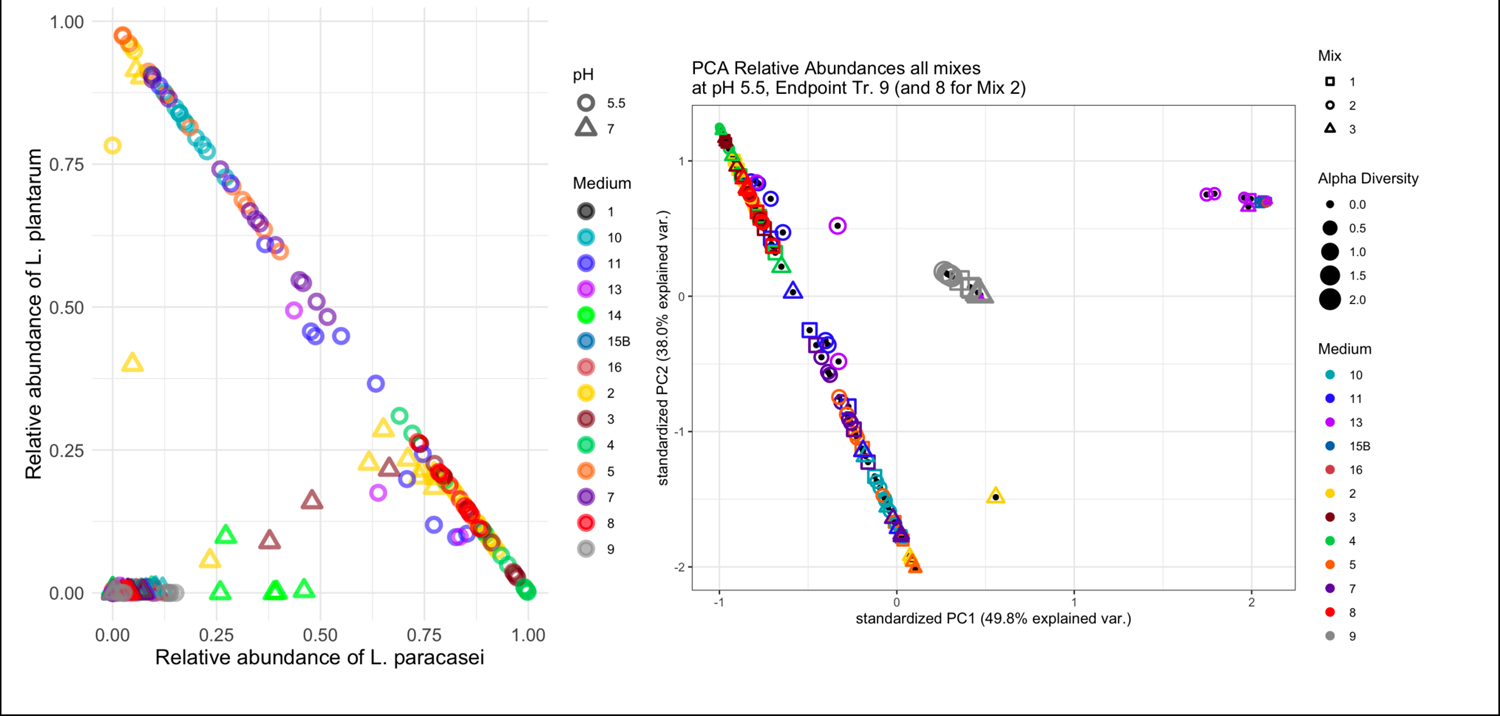
Left: Relative abundances of *L. paracasei* and *L. plantarum* in all the TrA1 experiment endpoints (including T8 for Mix-2), revealing their domination of the relative abundance profiles in pH 5.5. The strong negative correlation of these LAB was not considered an artefact of indistinguishable 16S, as these were different by 7 positions in respective V4 regions. Plus, if this strong negative correlation was an artefact of 16S rather than competitive exclusion, we would expect this artefact to be random and not medium-specific (as the relative domination of either species appears to be). Right: PCA of 16S relative abundance data for the endpoints of the TrA1 experiment, at pH 5.5. The strong diagonal is indicative of a negative correlation between *L. paracasei* and *L. plantarum* dominating relative abundance profiles. M9 poses an interesting exception, where medium type is the main driver of 16S variability and pH explains less, hinting at the buffering capacity of mucin in the absence of sugars. PC1 is defined by minimal media: M15B, M16, M13.

**Supplementary figure 5.**
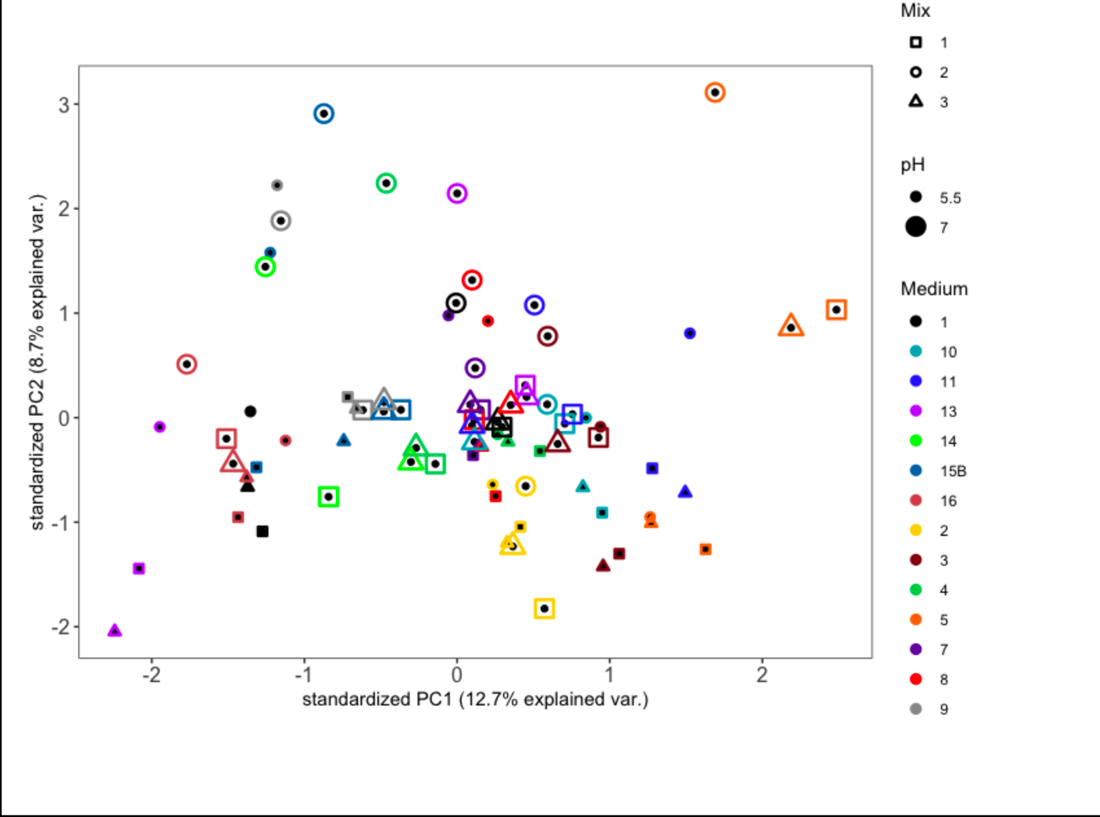
PCA of log2 fold changes (log2FC) of compounds with significant log2FC values (log2FC >|1|, FDR-corrected p < 0.05) relative to empty media controls. This provides a more specific focus on compounds that were significantly depleted or enriched by the communities supported in each medium. Most low-pH points cluster at the bottom half of the plot, whereas mix-2 points tend to cluster at the top half of the plot. This reveals the strength of the mix-dependent variation in logFC values of the metabolomics data, with mix-2 specifically having distinguishable profiles from their mix-1 and mix-3 counterparts along PC2 (per medium or medium * pH combinations). PC1 appears still to be defined by medium complexity, with at one extreme poor media (M16, M15B, M13), and at the other extreme richer media (M5, M11, etc.)

**Supplementary figure 6.**
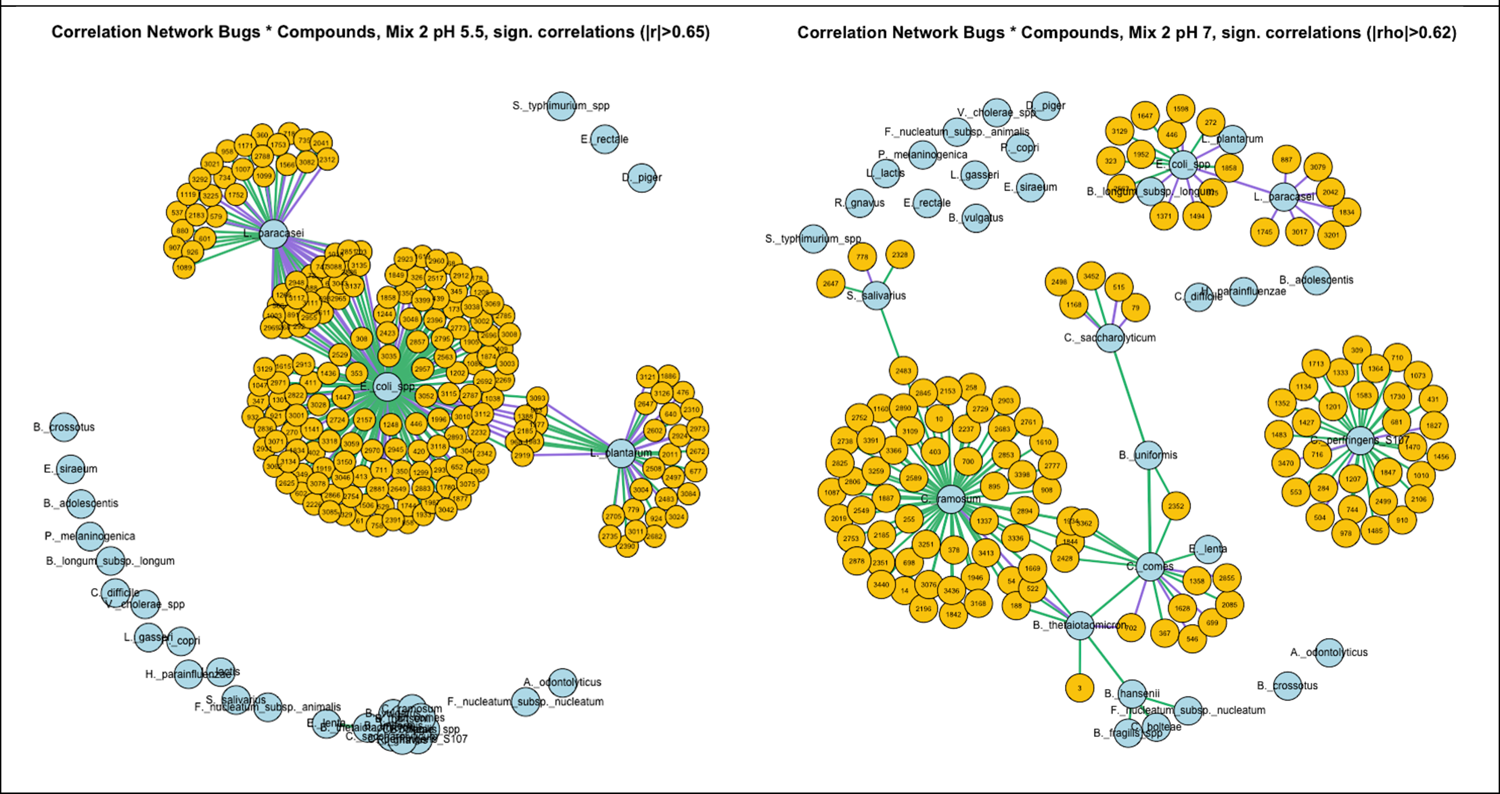
Correlation networks of 16S and metabolomics data of transfer 8 and 9 of Mix-2. Inter-ion correlations were removed for network visualisation purposes. Statistical significance of Spearman’s rank correlations was considered for p-values of <0.05 following correction for multiple inference using Holm’s method. Purple edges signify negative significant correlations, while green edges signify positive significant correlations. The first correlation network displays significant correlations (with Spearman rho > |0.65|) between species relative abundances and ions in as measured in pH 5.5, while the second network displays significant correlations (with Spearman rho > |0.62|) between species relative abundances and ions as measured in in pH 7.

**Supplementary figure 7.**
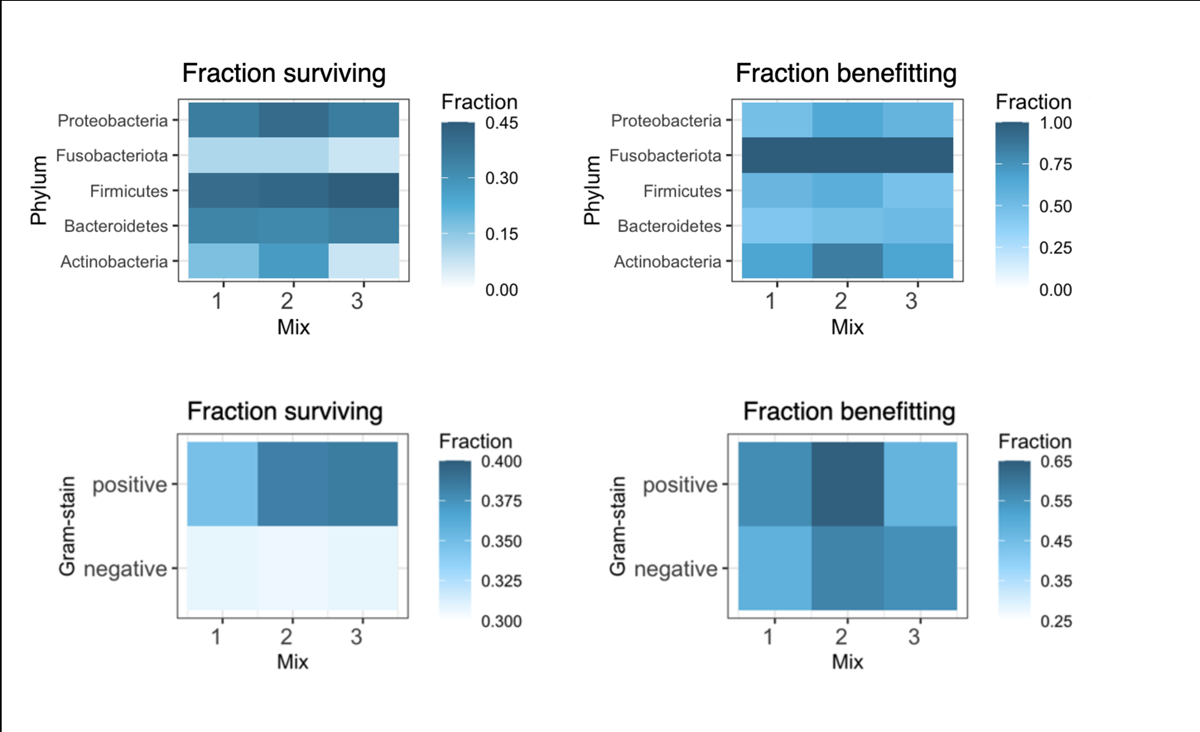
Heatmap of fractions surviving in co-culture and benefitting in co-culture per Phylum and Gram-stain. An expected pathway for benefitting from being in co-culture is through uptake, exchange or metabolism of extracellularly excreted compounds (i.e., cross-feeding), which might be influenced by the type of cell wall, specifically the number of cellular membranes through which compounds have to be transported. These heatmaps show that any strain’s propensity to survive, be an emergent survivor or experience emergent extinction appears associated not only with its phylum, but also with its cell wall architecture; Gram-positive strains (generally having a single membrane) display greater fractions of survival and of benefitting than Gram-negative strains (which have two membranes). Mix-2 was shown in Fig. 2d to have significantly higher metabolic enrichment, allowing cells being able to benefit from this enrichment to grow better; this may contextualise the highest fraction of benefitting being found for gram-positive strains in mix-2.

**Supplementary figure 8.**
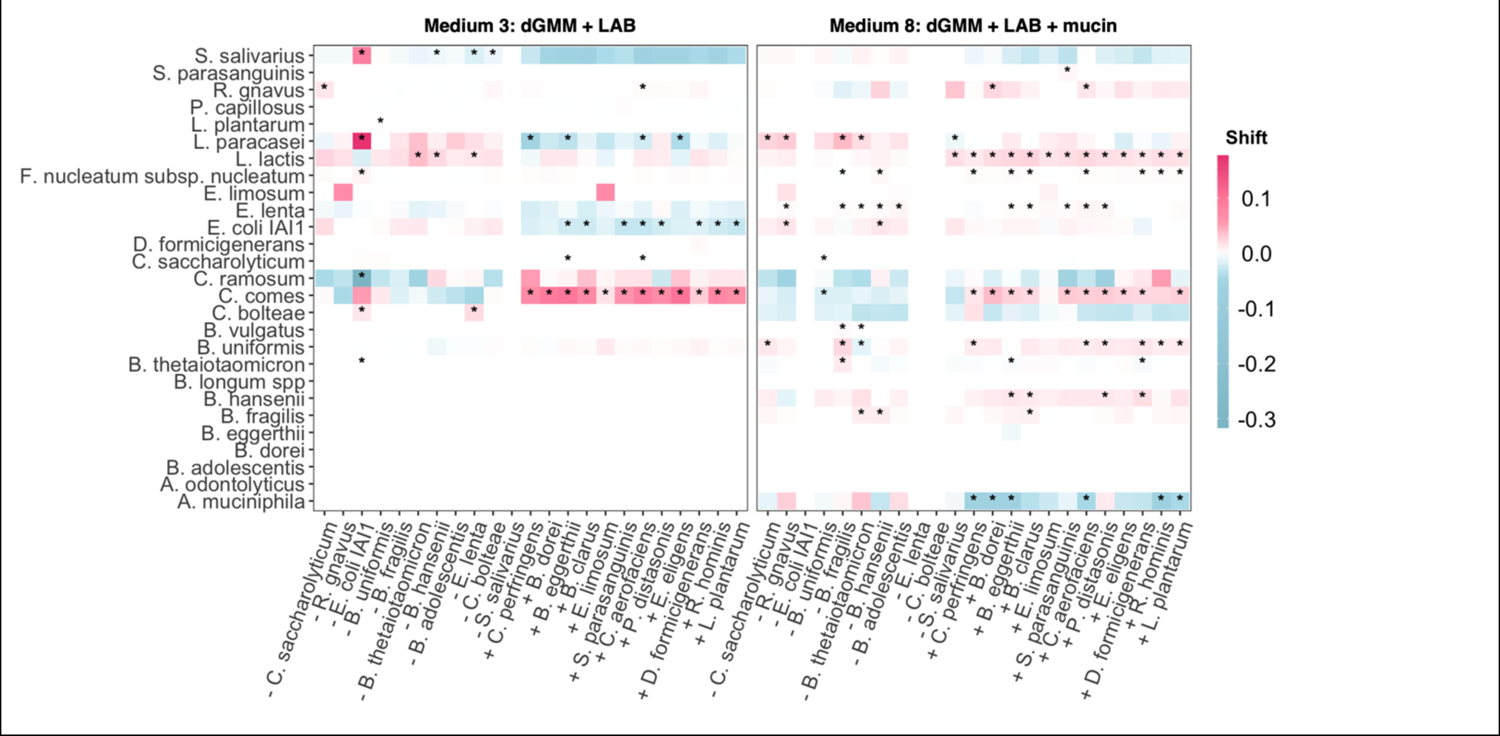
Heatmaps of shifts away from each species’ discounted abundance in the respective ‘All’ community (i.e., relative abundance with proportional redistribution of the singly excluded species, or the other singly added species), across all single exclusion and single addition experiments, separated by medium (M3 on the left, M8 on the right). This pair of heatmaps represent shifts in absolute values.

**Supplementary figure 9.**
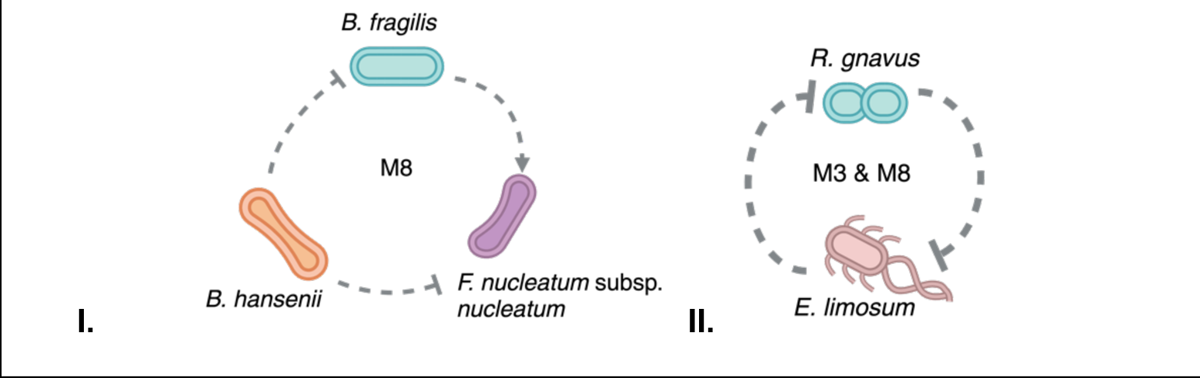
Inferred ecological landscapes. I. Interpretation of significant shifts observed for *F. nucleatum* subsp. *nucleatum* in M8. The single exclusion of *B. fragilis* lead to a significant downward shift for *F. nucleatum* subsp. *nucleatum*, implying a type of net positive effect of *B. fragilis.* Both *B. fragilis* and *F. nucleatum* subsp. *nucleatum* responded with a significant positive shift to the single exclusion of *B. hansenii*, implying a type of net negative effect of *B. hansenii* on these two (potentially pathogenically relevant) bacterial strains. The network implied, as displayed here, captures net interaction inferences, and could result from both direct as well as indirect interactions. For instance, the positive shift displayed by *F. nucleatum* subsp. *nucleatum* could be due to *B. hansenii* no longer inhibiting *B. fragilis;* an inferred ‘partner’ of *F. nucleatum* subsp. *nucleatum*. II. The single addition of *E. limosum* lead to the complete inhibition of *R. gnavus* – a bacterium that otherwise predominantly displays positive shifts across single additions. The inferred negative effect appears mutual when inspecting the effect of the single exclusion of *R. gnavus* on *E. limosum*: when *R. gnavus* was present in the community, *E. limosum* failed to achieve nonzero relative abundances, whereas when *R. ganvus* was singly excluded, *E. limosum* shifted up.

**Supplementary figure 10.**
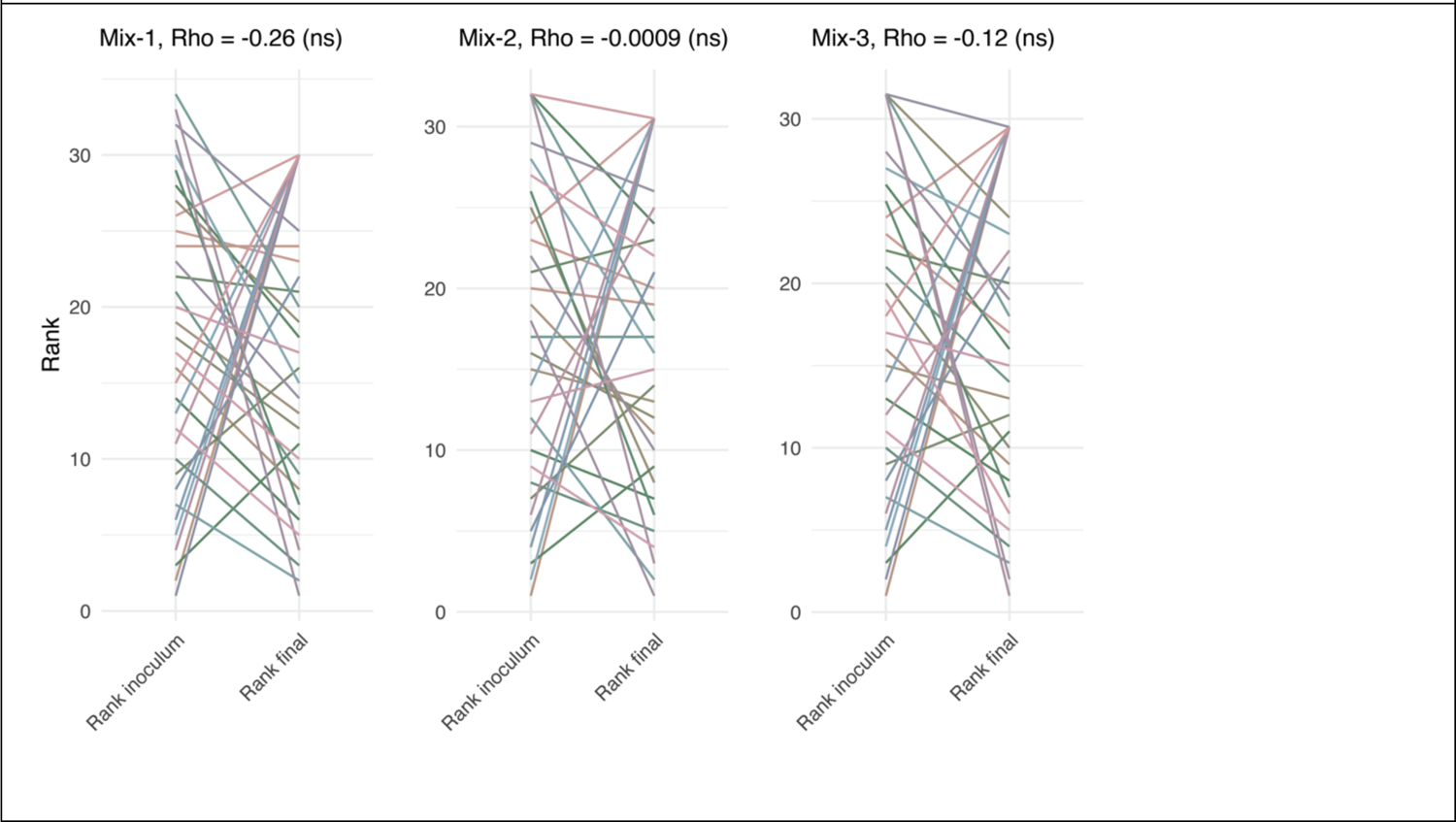
Analysis showcasing the negligible role of species’ inoculum load versus final load. For each mix, species’ mean relative abundances were ranked in respective inoculum and final (T9) community compositions. All spearman correlations were negative and insignificant, indicating that inoculum rank was unrelated to - and thus not predictive of - rank in final community composition.

**Supplementary figure 11.**
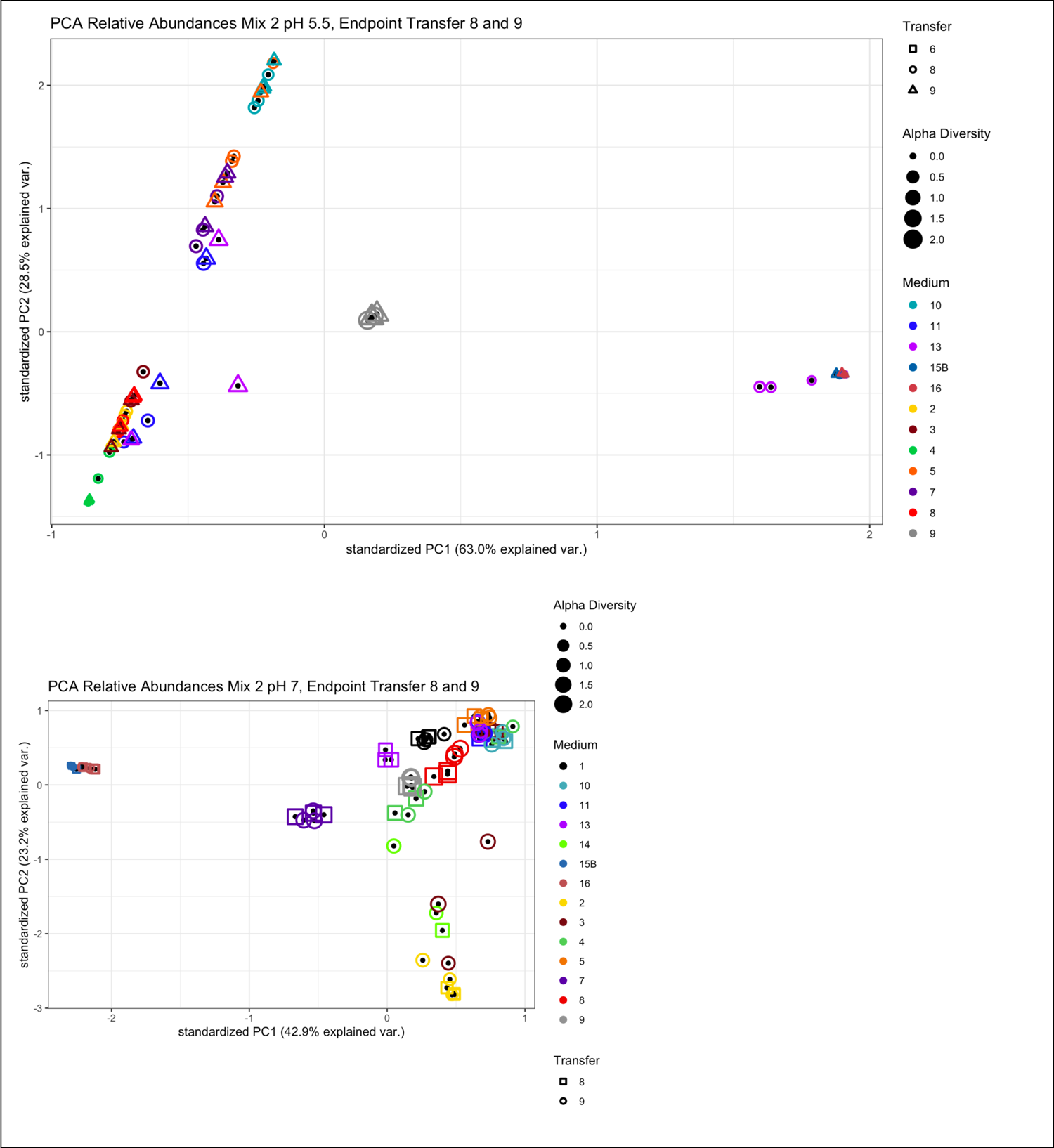
PCA of 16S relative abundance data of transfer 8 and 9, mix-2, showcasing that the difference between these timepoints is negligible. Mix- and or media-specific trends hold largely (except for M3, M13), and thus T8 data can be considered as supplementary endpoint data. For most media (if not all): inter-mix variation per timepoint > inter-time point variation per mix.

**Supplementary figure 12.**
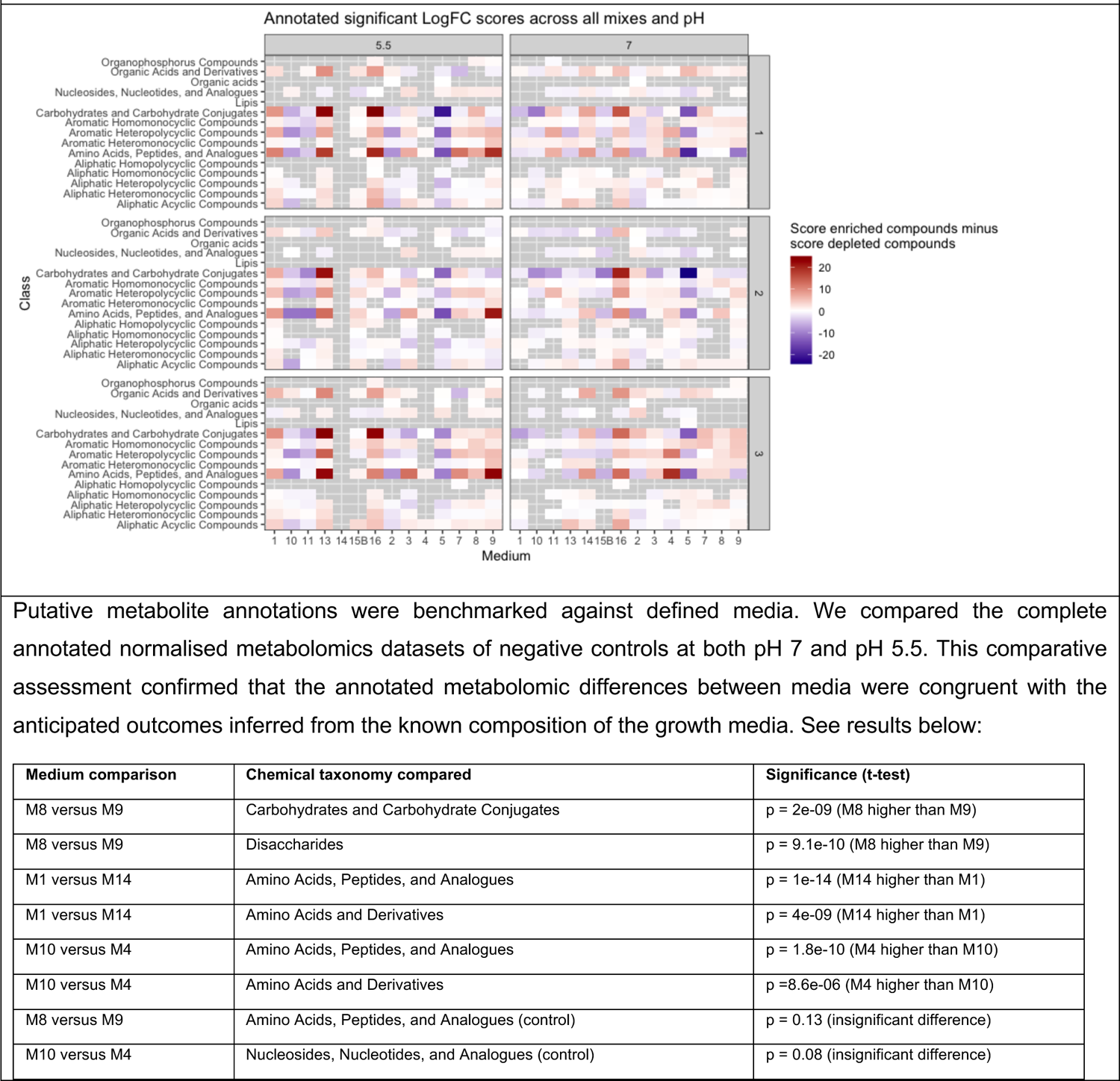
Annotated heatmap of LogFC metabolomics. Annotation of significantly enriched and depleted ions (logFC > |1|, FDR-corrected p-value < 0.05) across experimental conditions showcase pH-, mix- and medium-specificity. For Medium 9, amino acid depletion appears both mix-specific (with mix-2 in pH 7 showing lower depletion than mix-1 and 3, i.e., hinting at depletion due to pathogens present in initial mix) as well as pH-specific (with enrichment in pH 5.5 and depletion in pH 7). Despite medium 8 and 9 both being enriched with mucin, their relative depletion and enrichment profiles differ across many compound classes, once again hinting at the importance of simple sugars present in defining the community-level role of mucin. For some media and pH combinations, e.g., 1, 10, 11, 13 in pH 5.5, inoculation mix is little specific of depletion profiles.

**Supplementary Table 1:** Calculations of consistent community-dependent effects across all combinations of Species * Mix * Medium as found in the core experiment. Consistent community dependent effects (i.e., consistent across replicates within a 95% CI) were captured as ‘Status’, calculations of which can be found in the Methods section.

**Supplementary Table 2.** Spearman correlations (p < 0.05) between the normalised concentration of each compound across different growth media, and the media’s averaged Shannon alpha diversity (across all inoculation mixes at pH 7, core experiment). Of all 88 unique compounds (excluding buffer compounds) used in the 14 different growth media, 50 correlated significantly (p < 0.05) with mean alpha diversity, falling within 20 unique clusters after performing hierarchical clustering with height at 0.5 (i.e., several compounds have identical normalised concentrations across the different media and thus equal correlation strengths). All significant correlations were positive. Of all compound classes, ‘Others’, ‘Vitamins & Antioxidants’, ‘Salts and minerals’ and ‘Amino acids’ correlated most strongly with alpha diversity. None of the compounds in the class ‘Sugar’ correlated significantly with alpha diversity.

**Supplementary Table 3. A.** Overview of strains with nonzero abundances across replicates of starting inoculums for all three assembly experiments. Shaded in blue are strains that have nonzero abundances in the inoculums of all communities. Marked as red numbers are strains that are also classified as either a pathogen or probiotic, but were consistently present in either the inoculum and/or final compositions of probiotic- or pathogen specific mixes. **B.** Overview of co-culture survivalship of each species (i.e., ≥2/3 replicates with ≥ 0.5% relative abundance) in the core experiment (TrA1).

**Supplementary Table 4.** Spearman correlations between mean relative abundances *in vivo* and Shannon alpha diversity. Significance was adjusted using the Holm’s method. The average relative abundances of species were calculated from the 5.538 stool samples collected from healthy adult (age >= 18) included in curatedMetagenomicData (v 3.8.0; ^57^). *B. dorei* is emphasised in bold (due to its observed role, as captured in Fig. 5b). As expected, on average, a species contributes positively to alpha diversity (since every species with a nonzero abundance will by default contribute positively to alpha diversity, unless their abundance goes at the expense of the distribution of abundances of other community members). Note: one of the strongest correlators with alpha diversity *in vivo*, *C. comes*, also was one of the main survivors and benefiters in the core experiment (Fig. 4b), strongly benefitted from single additions (Fig. 5a); and displayed many correlations with metabolites (Suppl. fig. 6), emphasising the potential benefit *C. comes* gains from being in co-culture.

**Supplementary Table 5.** Overview of (experimental) species blanked in Fig. 5a & Supplementary Fig. 8.

